# Trifunctional sphinganine: a new tool to dissect sphingolipid function

**DOI:** 10.1101/2023.10.16.562528

**Authors:** Scotland Farley, Frank Stein, Per Haberkant, Fikadu G. Tafesse, Carsten Schultz

**Affiliations:** Oregon Health & Science University, Department of Chemical Physiology & Biochemistry, Portland, OR, 97239, USA; Department of Molecular Microbiology and Immunology, Portland, OR, 97239, USA; European Molecular Biology Laboratory, Proteomics Core Facility, 69117 Heidelberg, Germany

## Abstract

Functions of the sphingolipids sphingosine and sphinganine in cells are not well established. While some signaling roles for sphingosine have been elucidated, the closely related sphinganine has been described only insofar as it does not elicit many of the same signaling responses. The underlying mechanisms behind the cell biological differences between these lipids are not well understood. Here, we prepared multifunctionalized derivatives of the two lipid species that only differ in a single double bond of the carbon backbone. Using these novel probes, we were able to define their spatiotemporal distribution within cells. Furthermore, we used these tools to systematically map the protein interactomes of both lipids. The lipid-protein conjugates, prepared through photo-crosslinking in live cells and extraction via click chemistry to azide beads, revealed significant differences in the captured proteins, highlighting their distinct roles in various cellular processes. This work elucidates mechanistic differences between these critical lipids and sets the foundation for further studies on the functions of sphingosine and sphinganine.

## INTRODUCTION

Sphingolipids are an important and complex family of membrane lipids, including both structurally important lipids, such as sphingomyelin (SM) and complex glycosphingolipids^1^, and lipids involved in signaling cascades, including ceramide (Cer)^2^, sphingosine (Sph)^3^, and sphingosine-1-phosphate (S1P)^4^. While some roles for individual members of this family have been well-established, much remains to be uncovered about the structural requirements for carrying out specific biological functions, and the mechanisms by which sphingolipids exert their biological effects. A major gap in our understanding of the sphingolipid family exists around the two major sphingoid bases — sphingosine (Sph) and sphinganine (Spa)^5^. They are structurally extremely similar, differing only by one *trans* double bond, but have distinct metabolic sources and known functions. Spa is an early metabolite in *de novo* sphingolipid biosynthesis, and has long been thought of as an inert intermediate^1^. Spa is converted, through a series of enzymatic transformations, into SM, which can be hydrolized into Cer and then Sph.

The biological function of sphingosine is much better described than for sphinganine. Sph has well-established roles in promoting apoptosis^3^; exogenously applied sphingosine induces the cleavage of procaspase-3 and subsequent apoptosis^6^. It is still unclear exactly what intermediary pathways are involved, although the inhibition of MAPK^7^ and activation of sphingosine-dependent protein kinases^8, 9, 10^, have both been proposed. We also recently showed that sphingosine releases calcium from lysosomes when produced intracellularly, which begins a signaling cascade that ends with transcription factor EB (TFEB) translocation to the nucleus^11^. Critically, despite their structural similarity, Spa does not have these signaling properties when applied in the same way, neither the induction of apoptosis nor the release of calcium from lysosomes^3, 11^. In fact, the structural and signaling roles of sphinganine are elusive, as is the reason for the distinct differences in the cell biology of sphinganine and sphingosine.

To begin to unravel the cell biological differences between Spa and Sph, we synthesized trifunctional sphinganine to compare to the previously described trifunctional sphingosine^12^. Trifunctional lipids are designed to overcome several challenges in interrogating lipid biology: their fast metabolism, their weak and transient (but functionally important) interactions with proteins, and the difficulty in tagging them without disturbing the cellular location^13, 14^.

Trifunctional lipids add three functional groups to the endogenous lipid structure: 1) a terminal alkyne, for click chemistry with affinity tags or fluorophores, 2) a diazirine for photo-crosslinking with interacting proteins, and 3) a photocage to prevent premature metabolism and signaling^13^. Here, we investigated the metabolism and subcellular localization of both probes and showed that while the two sphingoid bases result in nearly identical pools of metabolites, the two lipids have subtle differences in their subcellular dynamics. Finally, we performed LC-MS/MS-based identification of probe-interacting proteins and described a network of lipid-interacting proteins with both distinct and overlapping interactions for each lipid base. These results provide structural insights into the differences and similarities in the biological roles for sphingoid bases.

## RESULTS

### Synthesis of trifunctional sphinganine

The initial steps of the synthesis of trifunctional sphinganine (TF-Spa, **1**) followed the previously described synthesis of trifunctional sphingosine (TF-Sph, **2**). The oxazolidine- protected ketosphingosine analog **3** was then subjected to a DIBAL reduction, which produced a mixture of 1-2 and 1-4 reduction products. The aliphatic ketone **4** was stereoselectively reduced to the alcohol, and the oxazolidine ring was opened under mild acidic conditions. The resulting bifunctional sphinganine derivative **5** was equipped with a caging group using coumarin chloroformate produced in situ from 7-diethylamino-4-hydroxymethylene-coumarin and phosgene (Scheme 1A) to give the trifunctional sphinganine (TF-Spa) derivative **1**.

**SCHEME 1.**
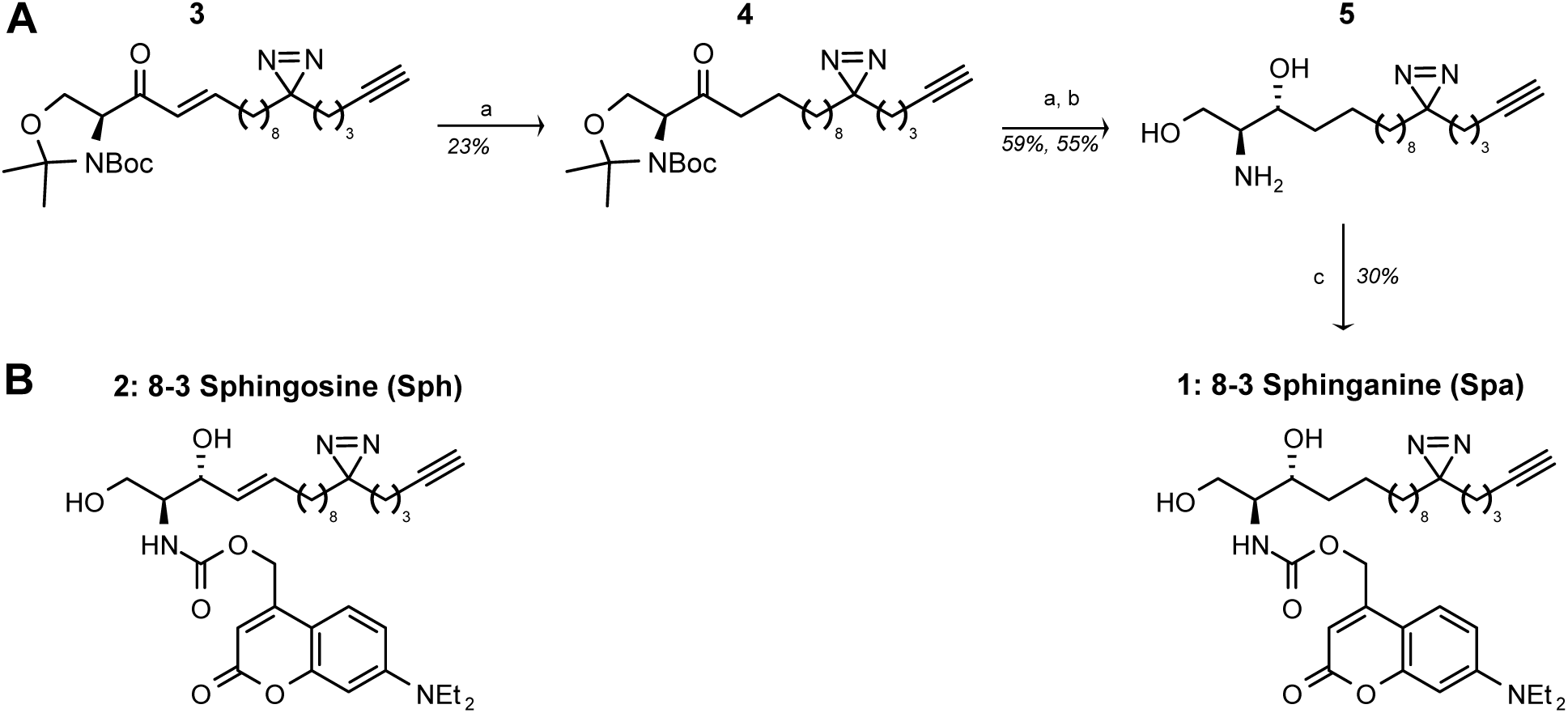
Synthesis of trifunctional sphinganine. (A) Synthesis of trifuntional sphinganine (**1**). (**a**) DIBAL-H, toluene, -78 °C; (**b**) HCl, MeOH; (**c**) 7-diethylamino 4-hydroxymethylene coumarin, phosgene; then DIEA, DCM, 0 °C. (B) Structure of trifunctional Sph (**2**), synthesized in parallel, with all characterizations matching previously reported NMR spectra^12, 18^.

### Biological fidelity of trifunctional sphingoid bases

A key consideration in any biological probe development effort is whether the derivatized probe faithfully recapitulates the properties of the endogenous molecule. A readily observable property of sphingosine is its ability to release calcium from lysosomes. This was described in detail using caged sphingosine to release the endogenous molecule with great temporal and spatial resolution^11^. In the same study, the authors showed that the release of sphinganine from its caged precursor, despite its structural similarity to sphingosine, was unable to produce a calcium spike. To verify that our trifunctional probes maintained the properties of their endogenous counterparts, we performed a similar uncaging experiment with TF-Sph and TF-Spa.

We treated live HeLa cells with 2 µm TF-Sph or TF-Sph, as well as the calcium-sensitive dye Fluo-4^15^. Using a dual-laser setup, we illuminated a region of a field of view with 375 nm blue light to uncage the sphingoid base, while imaging at 488 nm to observe the Fluo-4 fluorescence. As with the caged endogenous lipids, we observed an immediate increase in calcium fluorescence in the uncaged region in cells that had been treated with TF-Sph, but not in cells that had been treated with TF-Spa (representative images are shown in Fig 1A; normalized Fluo-4 fluorescence is shown in Fig 1B). Thus, we conclude that the diazirine and terminal alkyne are not significant enough modifications to perturb the biological function of the lipid.

**FIGURE 1.**
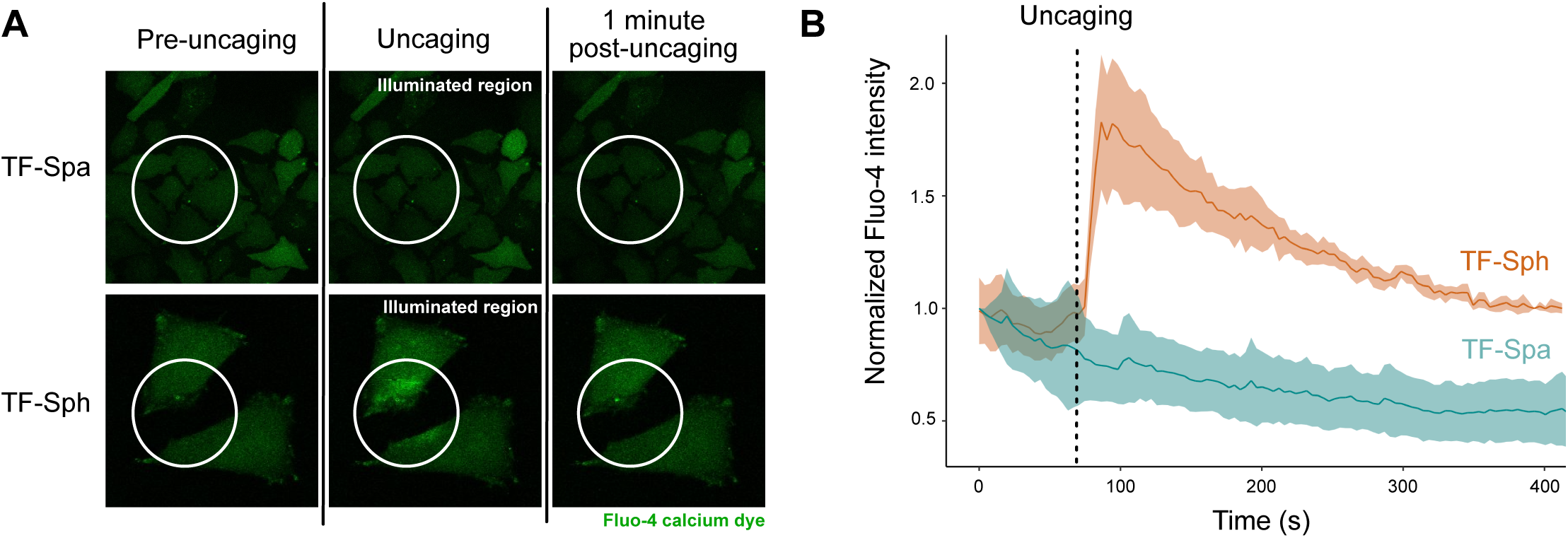
Biological fidelity of trifunctional sphingoid bases. HeLa cells treated with either TF-Spa or TF-Sph, as well as the calcium-sensitive dye Fluo-4, were subjected to uncaging while imaging Fluo-4 fluorescence. **(A)** Representative images of cells before, immediately after, and 1 min after uncaging. **(B)** Fluo-4 fluorescence intensity, normalized to initial intensity reading for each cell, for 15 cells (TF-Sph), or 18 cells (TF-Spa), over two independent experiments. The error band is standard deviation; the measure of center for the error band is the mean.

### Metabolism of novel lipid derivatives

A further test of the biological fidelity of the trifunctional probes is whether they are recognized by the endogenous biosynthetic machinery. Sphinganine is nominally upstream of sphingosine in the sphingolipid metabolic pathway, but both molecules are substrates of ceramide synthase, and may be expected to be incorporated into similar lipid classes, in particular ceramide (Cer), sphingomyelin (SM), and, via the salvage pathway, phosphatidylcholine (PC)^1^ (Fig. 2A). To investigate the metabolism of TF-Spa and TF-Sph, we treated Huh7 cells with the trifunctional lipid derivatives, extracted the lipids after 60 min, labeled the lipid extract fluorescently with Alexa555-azide via click chemistry and performed TLC (Fig. 2B). We turned to Huh7 cells because they are a frequently used model liver cell line for various diseases including viral infections^16, 17^. In addition, this added a dataset that was broadly similar to the previously published metabolism of bifunctional Sph^18^. In line with what has previously been shown with radiolabeled sphingosine^19^ and bifunctional Sph in mouse embryo fibroblasts^18^, the majority of the sphingoid base probes were incorporated into SM and PC, with some appearing as Cer (Fig 2C).

**FIGURE 2.**
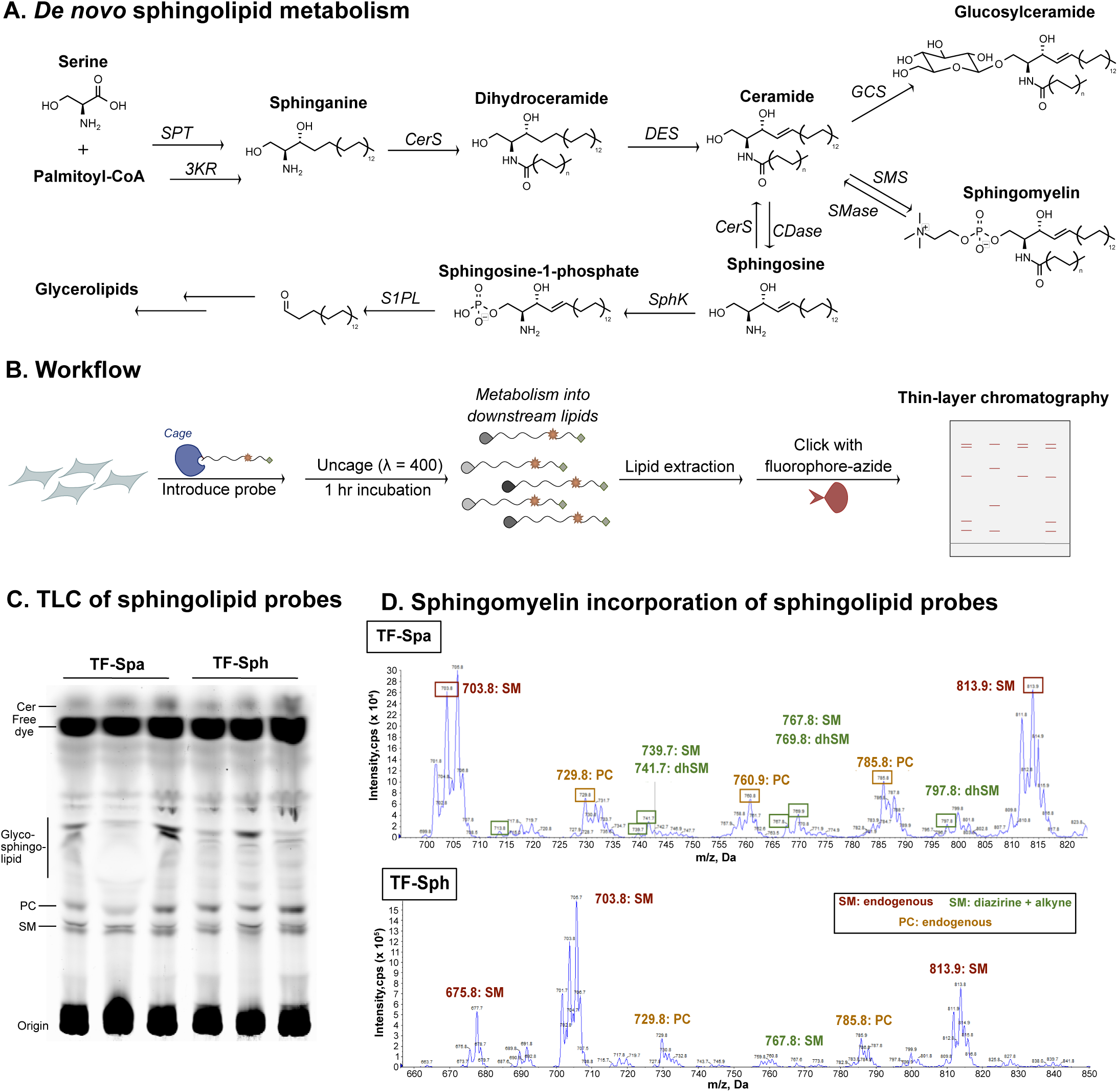
Metabolism of TF-Spa and TF-Sph. (**A**) Workflow. Cells are treated with the lipid of interest, exposed to 400nm light to uncage the probe, which is allowed to metabolize for 1 h prior to lipid extraction, click reaction with a fluorescent azide, and separation by thin-layer chromatography. (**B**) Overview of sphingolipid metabolism. Abbreviations: SPT = serine palmitoyl transferase; CerS = ceramide synthase; DES = ceramide desaturase; GCS = glucosylceramide synthase; SMS = sphingomyelin synthase; SMase = sphingomyelinase; CerAse = ceramidase; SK = sphingosine kinase; S1PL = sphingosine-1-phosphate lyase (**C**) TLC of TF-Spa and TF-Sph 1 h after uncaging, in Huh7 cells. Three biological replicates are shown for each probe. (**D**) Precursor ion scanning for precursors of phosphocholine in extracts of Huh7 treated with each sphingoid base, uncaged, allowed to metabolize for 1 h prior to lipid extraction and injection into the LC- MS/MS.

To further verify that both TF-Spa and TF-Sph are incorporated into SM, we performed targeted LC-MS/MS on lipid extracts prepared as described above, using precursor ion scanning to identify precursors of phosphocholine at the elution time for SM (LC-MS/MS methods are described in detail in Supplemental Information). We observed both the expected endogenous precursors of phosphocholine: SM (d18:1/16:0 and d18:1/24:1) and PC (34:1 and 36:2). We also observed SM with masses for the diazirine/alkyne modified probe backbone. TF- Spa appeared in a wider variety of SM species: both the SM and dihydroSM (dhSM) species with a 16:0 fatty acid; both the SM and dhSM species with an 18:0 fatty acid; and the dhSM species with a 20:0 fatty acid. TF-Sph appeared in the SM species with an 18:0 fatty acid (Fig. 2D).

### Subcellular localization of novel lipid probes

Next, we wanted to examine the subcellular localization of each sphingoid base over time. Previous work with bifunctional sphingosine in sphingosine 1-phosphate lyase knockout mouse embryo fibroblasts (MEFs) indicated that sphingosine initially localized to lysosomes, subsequently moved to the ER and later to the Golgi apparatus^18^. We performed a time course experiment in the liver-derived Huh7 cells, crosslinked each probe either 5, 30, or 60 minutes after uncaging, and co-stained with various organelle markers, including LAMP1 for lysosomes, PDI for the ER, and giantin for the Golgi (Fig. 3A and 3B). We calculated Pearson’s coefficients as a measure of co-localization between the organelle marker and the lipid signal for each cell, using a Cellprofiler pipeline (Fig. 3C).

**FIGURE 3.**
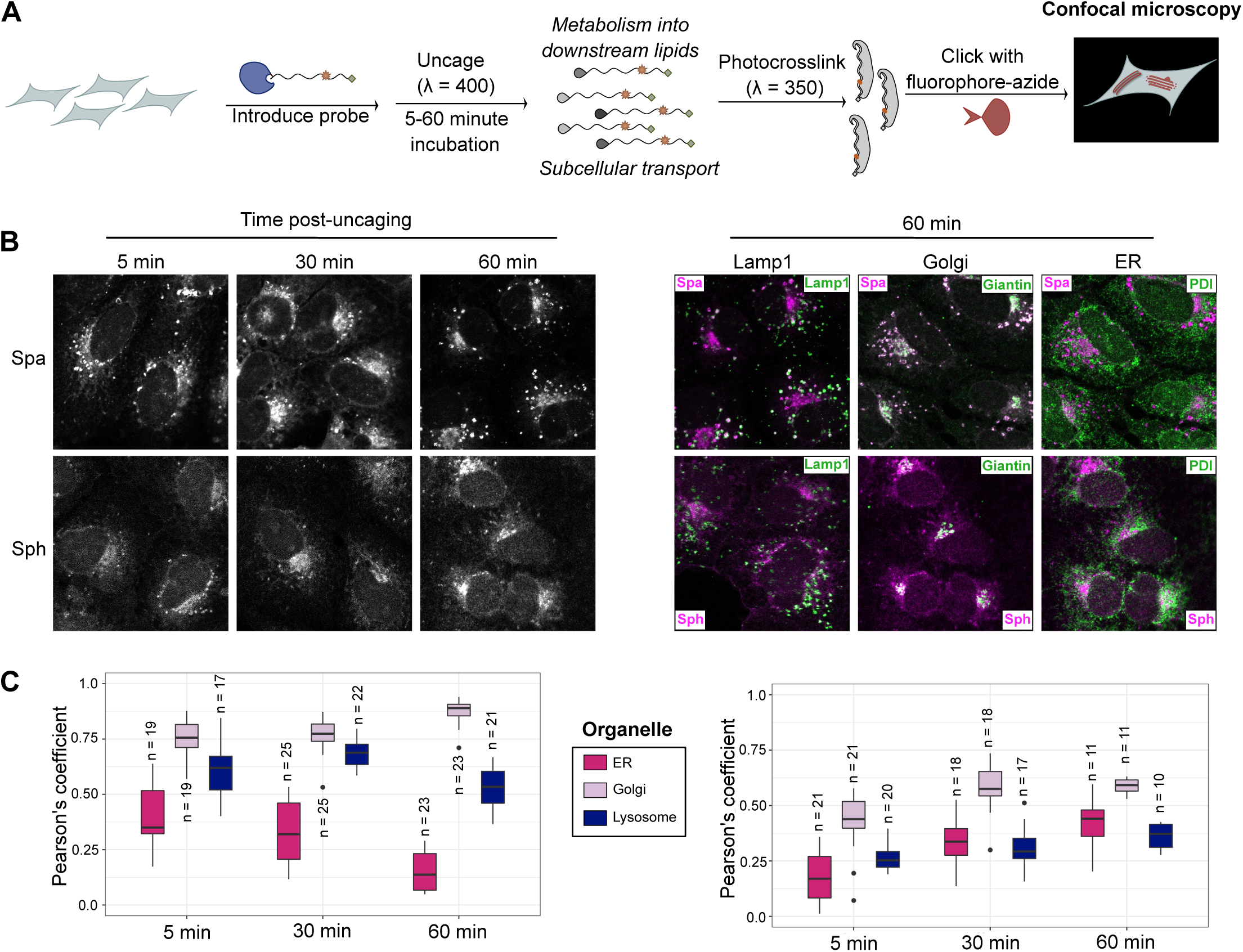
Subcellular localization of lipid probes. (**A**) Experimental design. Huh7 cells were treated with TF- Spa or TF-Sph, exposed to 400nm light to uncage the probe, and allowed to metabolize the probe for 5, 30, or 60 min. Lipid-protein complexes were photo-crosslinked with 350nm light, fixed, subjected to click reactions with a fluorescent azide, immune-stained for organelle markers, and visualized by confocal microscopy. (**B**) Representative images of each lipid at each timepoint; images are representative of four biological replicates over two independent experiments. (**C**) Colocalization of each lipid probe with markers for organelles; PDI for the ER, Giantin for the Golgi, and Lamp1 for the lysosome. Pearson’s correlation coefficients were calculated between the organelle marker and the lipid for individual cells using a Cellprofiler pipeline. Number of cells used to calculate correlation coefficients are indicated above their respective boxplots.

Both sphingoid bases were highly enriched in lysosomes and the Golgi; TF-Spa began at 5 minutes with relatively high association with the ER, which decreased over time in favor of the Golgi; this reflects what is known about de novo sphingolipid biosynthesis, where sphinganine synthesized in the ER is converted into Cer and then trafficked to the Golgi. This trend was less dramatic with TF-Sph, where the co-localization with both the ER and Golgi gradually increased over time. These observations generally underline what is known about both de novo and salvage sphingolipid biosynthesis and localization, and position these tools well to interrogate systems where these homeostatic states are perturbed.

### Identification of interacting proteins of TF-Spa and TF-Sph

With the metabolism and localization of each probe established, we next sought to use the photo-crosslinking function of the trifunctional lipid derivatives to compare the protein interactome of each lipid backbone. Such interactomes have been described for cholesterol^20^, bifunctional sphingosine^18^, bifunctional fatty acid^21^, trifunctional sphingosine, FA and DAG^12^, and trifunctional PI^22^, PI(3,4)P_2_, and PI(3,4,5)P_3_^23^. We treated Huh7 cells with each of the sphingoid base derivatives in two biological replicates, and also included two biological replicates for each probe that had not been exposed to 350nm light for crosslinking (-UV) as a negative control. We photo-crosslinked the sphingoid bases 15 min after uncaging, to allow for some metabolism but before sphingosine-1-phosphate lyase was able to degrade the sphingoid backbone. We used azide-agarose beads to enrich proteins that had been crosslinked to the alkyne-bearing lipids and digested off the isolated proteins from the beads with trypsin, as previously described^23^. Peptides were isotopically labeled using the TMT-16 platform for multiplexing and analyzed by LC-MS/MS (Fig 4A). Raw signal intensities for each channel were normalized based on variance stabilization. 1129 proteins were identified. To assess whether there was appreciable enrichment of proteins in the +UV condition over the -UV condition, the ratio of the intensity of each protein in the +UV condition verses the -UV condition was calculated; the log_2_ of this ratio is displayed in Fig 4B, highlighting the enrichment of distinct subsets of proteins in each probe condition, and good overall enrichment of proteins for the irradiated conditions.

**FIGURE 4.**
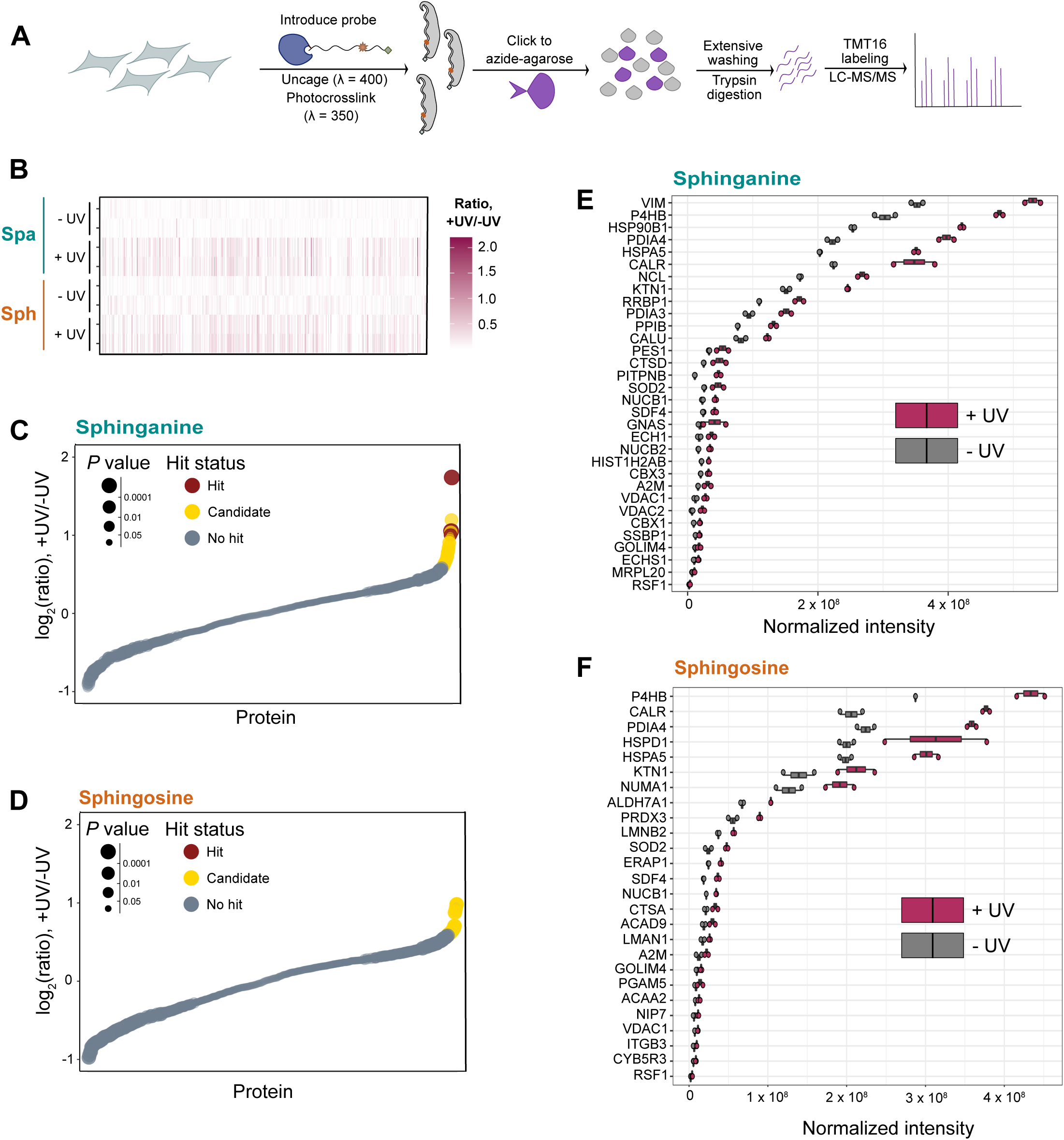
Identification of lipid-binding proteins (**A**) Experimental design. Cells are treated with the probe of interest, illuminate to uncage and then photocrosslink the probe, and lysed by sonication. Lysates are subjected to a click reaction with azide agarose, and beads are extensively washed to remove non-covalently bound proteins. Bound proteins are trypsinized off the beads, desalted and TMT-labeled for quantitative proteomics, and analyzed by LC-MS/MS. (**B**) Heat map of intensity ratios for 1129 proteins identified in the +UV illuminated samples over the -UV samples for each probe. (**C-D**) Overview of identified hit proteins for each probe, by Limma analysis. “Hits” are defined as proteins with a false discovery rate less than 0.05 and a fold change of at least 2-fold in the +UV over the -UV. “Candidates” are defined as proteins with a false discovery rate less than 0.2 and a fold change of at least 1.5-fold. (**E-F**) Normalized intensities of proteins identified as hits or candidates by the Limma analysis for (**E**) TF-Spa; (**F**) TF-Sph

To identify the most robust interacting proteins for each probe, the normalized signal intensities were subjected to Limma analysis to calculate fold changes and *p*-values of the intensity in the +UV over the -UV samples (Fig 4C and 4D). Hits and candidates were selected based on fold change and false discovery rate criteria: “hit” proteins are those with a fold change greater than 2 and a false discovery rate less than 0.05; “candidate” proteins are those with a fold change greater than 1.5 and a false discovery rate less than 0.2. We identified 4 hits and 28 candidates for TF-Spa and 26 candidates for TF-Sph. The normalized signal intensities for all hits and candidates are displayed in Fig 4E-F, for each probe. A range of proteins were identified for each probe, including many known lipid binding proteins. TF-Spa and TF-Sph both pulled down proteins already shown to interact with bifunctional sphingolipids, including VDAC1/2 (shown to bind bifunctional ceramide^24^), and cathepsin D (previously described to bind ceramide^25^, and pulled down in a study of bifunctional sphingosine^18^).

To functionally compare the hits and candidates for each probe, the GO terms for each protein were analyzed (Fig 5A, Fig 5B). Not unexpectedly, for both probes, there was an enrichment in proteins involved in lipid transport. A large number of homeostatic proteins were identified, including many ER-resident proteins involved in protein processing. Both probes interacted with components of cell distress pathways, including ER proteins involved in the unfolded protein response and factors contributing to intrinsic apoptosis (CTSD, VDAC1/2).

**FIGURE 5.**
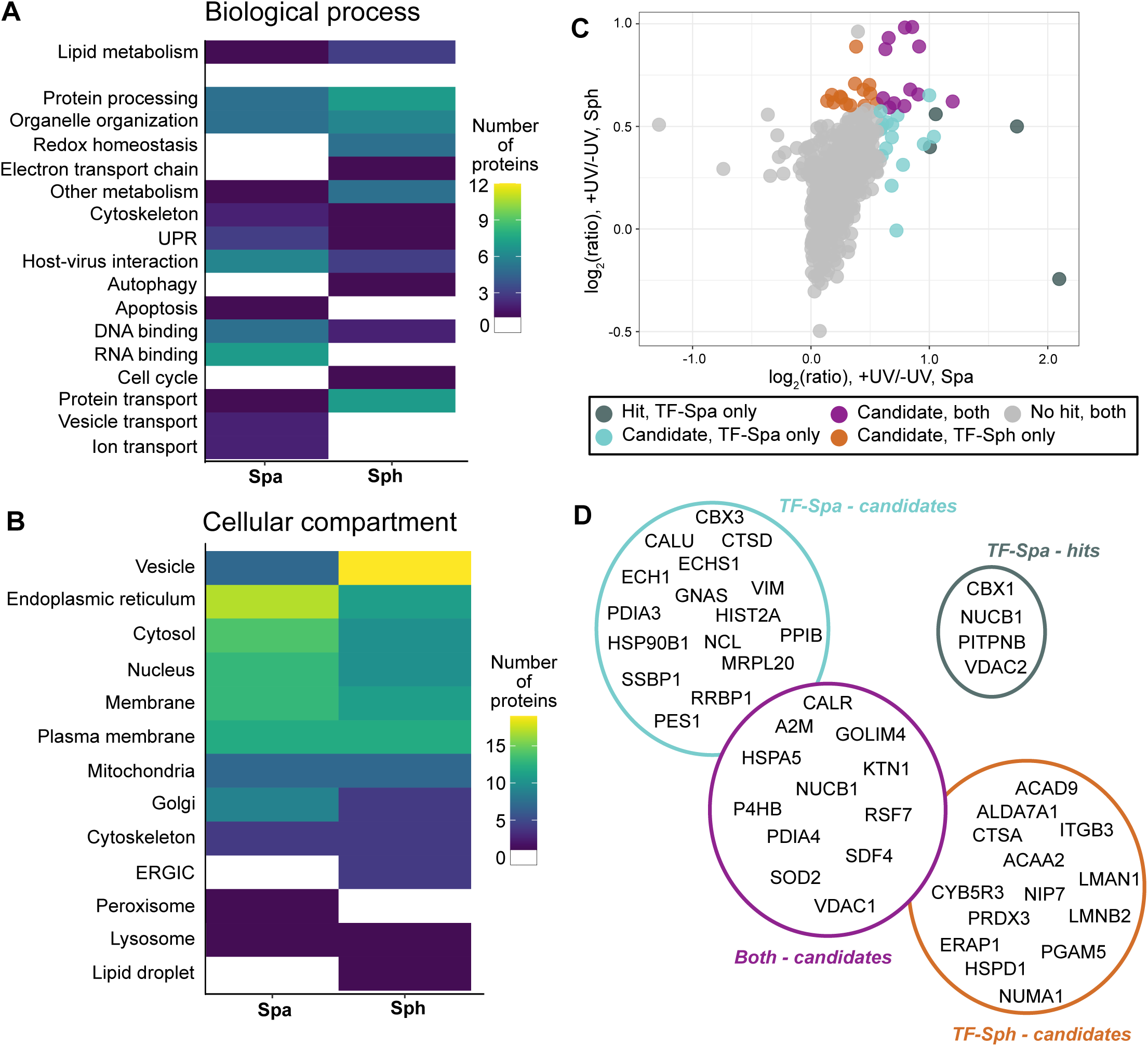
Analysis of lipid-binding proteins (**A**) GO-term enrichment of proteins identified as hits or candidates for each probe. The number of probe-binding proteins involved in each biological process for each probe is shown. (**B**) GO-term enrichment of proteins identified as hits or candidates for each probe. The number of probe-binding proteins found in each cellular compartment for each probe is shown. (**C**) Comparison of TF-Spa and TF-Sph interacting proteins. The log_2_(FC) for the UV condition over the -UV condition for each probe is shown on each axis. “Hit” and “candidate” categories are defined as described above. (**D**) Map of overlap of lipid-binding proteins, illustrating which proteins belong in the categories delineated in (C)

Congruous with our previous microscopy findings, proteins were found annotated to various compartments within the cell, especially the mitochondria, the ER, and the nuclear envelope; TF-Spa had the highest number of ER-associated proteins while TF-Sph had the highest number of vesicle-associated proteins.

Finally, we analyzed the overlap between the hits and candidates observed for each probe. TF-Sph and TF-Spa both had a number of unique hits/candidates (20 for TF-Spa and 12 for TF-Sph), and shared twelve candidates (Fig 5C, breakdown of which proteins are overlapping in Fig 5D). While it is not surprising that there is some overlap in the protein associations of these two molecules, the divergence of the pulled-down proteins between the two sphingoid bases is striking, given their structural similarity, suggesting that at least some of their differences in biological function may be derived from different interactions with the cellular proteome.

## DISCUSSION

In this work, we present the synthesis of a novel multifunctional sphinganine probes to compare the interactions of two basic sphingoid bases, sphingosine and sphinganine. The two sphingoid base derivatives (TF-Spa and TF-Sph) are chemically and structurally extremely similar. However, they are clearly distinguished in their signaling properties, metabolic connections, and protein interactions, and are functionally very different. While more is known about the functions of sphingosine, sphinganine has often been compared to sphingosine and found to be unable to elicit the same signaling responses as sphingosine. There are several hypotheses for these differences, but they have not been extensively tested. Potentially, the signaling properties of these molecules are controlled largely by their controlled subcellular metabolic release — sphinganine, produced early in sphingolipid biosynthesis in the ER and rapidly transformed into dihydroceramide^1^, would only be present in large amounts under extremely unusual circumstances. It is not well described where in the cell this molecule might be located absent the strictures of endogenous *de novo* synthesis. Sphingosine, meanwhile, is produced by the action of ceramidases which are located in many compartments of the cell, including the lysosome (acid ceramidase), the plasma membrane (neutral ceramidase), and the ER (alkaline ceramidase), usually in response to stress stimuli^26^. Alternatively (or in addition), the slight structural difference between the structure of these two lipids may be sufficient to distinguish their respective affinities for proteins, and thus their influence on signaling pathways in the cell. It might be speculated that the double bond in Sph changes the pKa of the vicinal secondary hydroxy group, leading to a different molecular interaction potential.

We showed that TF-Spa and TF-Sph were incorporated very similarly into the sphingolipid biosynthetic pathway, using thin-layer chromatography of the click-labeled lipid extracts and LC-MS/MS to identify metabolites with the unnatural probe backbone, at least at the time points covered. We further showed the two probes recapitulate the distinct phenotypes of the endogenous lipids with respect to calcium signaling: TF-Sph uncaging induces an immediate spike of calcium from live cells, while TF-Spa uncaging does not. We showed that both sphingoid bases accumulated in the lysosome, and over time, in the Golgi. In the absence of a functional probe for sphinganine, such a time course of subcellular localization of exogenously applied sphinganine had never before been described. The similarity of localization between TF-Spa and TF-Sph suggests that the differences in their biological effects is not driven primarily by differences in location — especially since many studies that distinguish between Spa and Sph in calcium signaling and apoptosis were performed using exogenously applied lipids^6, 11^.

The profiling of protein interacting partners was based on -UV control conditions to select genuine probe-binding partners. We identified several proteins that overlapped with previous findings of bifunctional sphingosine and ceramide, including cathepsin D and VDAC1/2. We observed some overlap in the hits from TF-Spa and TF-Sph; many of these were abundant ER proteins (eg, P4HB, PDIA4). Interestingly, Sph also pulled down cathepsin A, which, while functionally related to cathepsin D, has not been shown to depend upon lipids for its function the same way as cathepsin D. TF-Spa interacted with more nuclear proteins and nuclear envelope proteins, while TF-Sph interacted with more vesicular proteins and proteins involved in subcellular transport. TF-Sph and TF-Spa had 19 and 20 unique hits, respectively, and shared 13 hits. The unexpected high number of unique hits for such structurally similar lipids suggests that the *trans-*double bond is sufficient to drive a certain amount of discrimination between protein interactions of the two lipids. Notably, while our microscopy experiments show both lipids occupying relatively similar compartments in the cell, interacting proteins were found from overlapping but non-identical regions of the cell, with TF-Spa pulling down a higher number of ER- and Golgi-associated proteins, while TF-Sph pulled down a striking number of vesicle-associated proteins. Future studies will interrogate the interactions described here for functional significance and relevance in known sphingolipid signaling cascades. It is likely that some cellular functions of Spa and Sph, respectively, will depend on the proteins that turned up as unique hits. Therefore, the proteomic results will serve as a good starting point to study the functions of both sphingoid lipids in the future.

## ACKNOWLEDGMENTS

Targeted LC-MS/MS of sphingomyelin and phosphocholine (in Figure 2) was performed with the help of the Pharmacokinetics Core at OHSU, run by Andrea DeBarber. This work was supported by a funding from the National Institutes of Health (NIAID R01 AI141549-02, to FGT and CS).

## AUTHOR CONTRIBUTIONS

S. E. F. and C. S. designed the study; S. E. F. designed and executed the chemical synthesis and performed cell-based experiments; P. H. and F. S. ran and analyzed the proteomics samples; S. E. F. wrote the original draft of the manuscript; C. S. edited the manuscript; all authors reviewed the manuscript; C. S. supervised and and administered the project; F. G. T. and C. S. acquired funding.

## METHODS

### Materials

#### Cell culture

Huh7 cells, obtained from ATCC, were maintained in standard tissue-culture treated vessels in DMEM supplemented with 10% FBS, 1% nonessential amino acids and 1% penicillin-streptomycin at 37 °C and 5% CO_2_.

#### Chemicals and reagents for synthesis

All chemicals were purchased from commercial suppliers and were used without further purification. Solvents were of ACS chemical grade (Fisher Scientific) and were used without further purification. Commercially available starting reagents were used without further purification. Analytical thin-layer chromatography was performed on silica gel 60 F254 aluminum-backed plates (Millipore Sigma), and spots were visualized either by UV light (254 nm or 365 nm), or potassium permanganate staining (1.5 g KMnO_4_, 10g K_2_CO_3_, and 1.25 mL 10% NaOH in 200 mL of water). Flash column chromatography was performed with manually packed columns using Thermo Scientific Chemical Silica gel (0.035-0.070mm, 60 Å). High pressure liquid chromatography (HPLC) was performed on a Varian Prostar 210 (Agilent) using Polars 5 C18-A columns (Analytical: 150 x 4.6 mm, 3 µm; Preparative: 150 x 21.2 mm, 5 µm). Mass spectra were obtained on a Advion Expressions CMS mass spectrometer. ^1^H NMR spectra were recorded on a Bruker DPX spectrometer at 400 MHz.

Chemical shifts are reported as parts per million (ppm) relative to solvent references. Spectral characterization can be found in the Supplemental Information.

### Methods

#### Synthesis of TF-Spa (**1**)

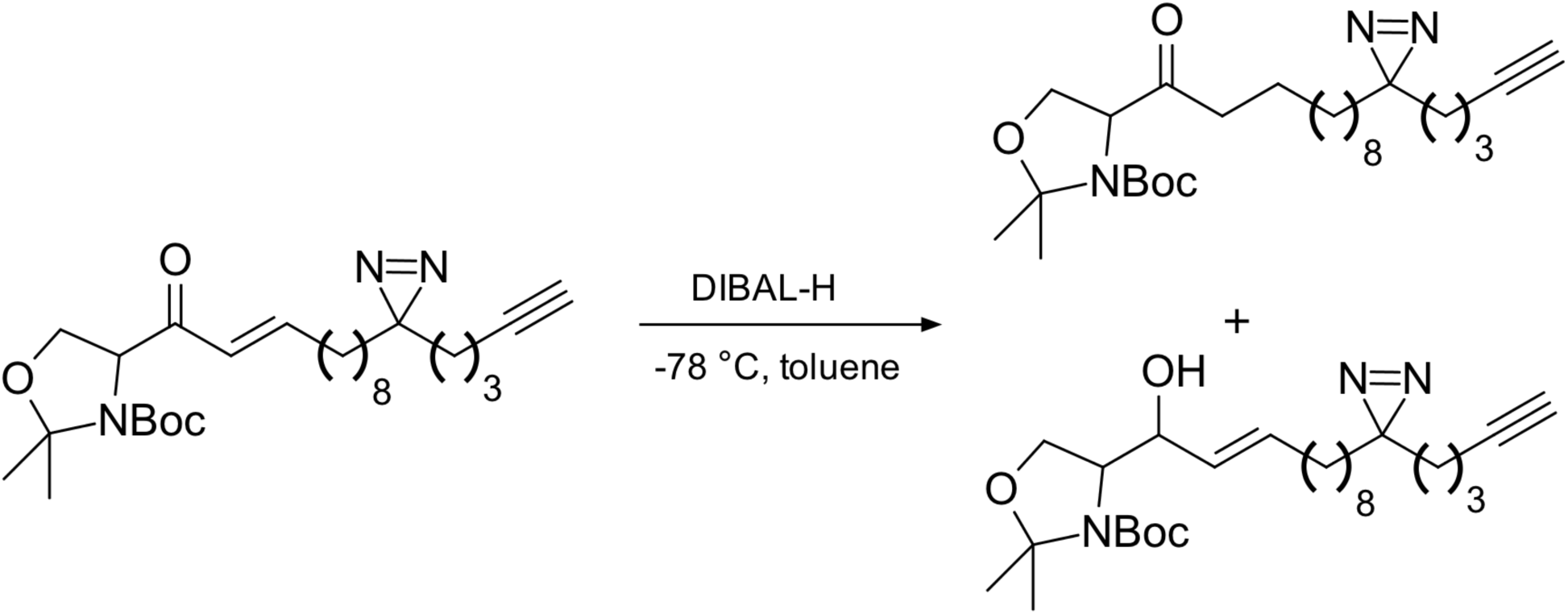

**Tert-butyl 2,2-dimethyl-4-{11-[pent-4-yn-1-yl)-3H-diazirin-3-yl]undecanoyl}-1,3-oxazolidine-3- carboxylate (4)** Compound **3** (0.50 g, 1.056 mmol, 1 equivalent) was dissolved in anhydrous toluene (15 mL) and cooled to -78 °C. DIBAL-H (1.1 mL, 1.056 mmol, 1 equivalent) was added, dropwise. After ∼1 minute, took a TLC (4:1 hexanes:ethyl acetate, potassium permanganate visualization), which indicated the production of two products: the 1,4 reduction product, the ketone (**4**), with an Rf of 0.7 (Rf of **3** in this system is 0.65); and the 1,2 reduction product, the alpha-beta-unsaturated alcohol, with an Rf of 0.3. The 1,4 and the 1,2 reduction products were separated by flash chromatography (4:1 hexanes:ethyl acetate), producing ketone **4** as a pale yellow oil (0.109 g, 23 % yield), as well as the alpha-beta-unsaturated alcohol (0.100 g, 20% yield). ^1^H NMR (400 MHz, CDCl_3_) δ = 4.472 (dd, J = 7.4, J = 3.2, 1H), δ = 4.325 (dd, J = 7.6, J = 2.8, 1H), δ = 4.155 (m, 2H), δ = 3.915 (m, 2H), δ = 2.494 (q, J = 8, 2H), δ = 2.178 (td, J = 6.8, J = 2.8, 2H), δ = 1.962 (t, J = 1.4, 1H), δ = 1.725 (s, 2H), δ = 1.666 (s, 2H), δ = 1.586 (s, 4H), δ = 1.513 (m, 9H), δ = 1.424 (s, 4H), δ = = 1.365 (m, 6H), δ = 1.25 (m, 12H), δ = 1.090 (m, 2H); ^13^C NMR (400 MHz, CDCl_3_) δ = 208.801, 208.360, 207.014, 152.485, 151.531, 138.083, 115.127, 95.247, 94.547, 83.568, 80.999, 80.663, 68.993, 68.310, 65.844, 65.486, 65.287, 39.184, 38.537, 33.029, 32.427, 31.947, 31.051, 29.819, 29.510, 29.430, 29.371, 28.557, 28.475, 26.324, 25.512, 24.967, 23.933, 23.205, 22.873, 18.077;

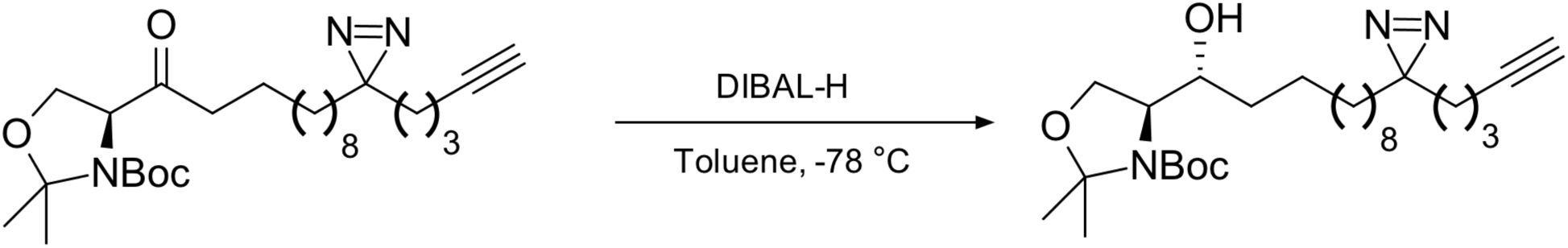

[M + H]+, calcd. for C_29_H_39_N_3_O_4_, 476.3490, found 476.5.

**(1R)-1-[3S)-5,5,-dimethyloxolan-3-yl]-11-[3-(pent-4-yn-1yl)-3H-diazirin-3-yl]undecan-1-ol (17)** Compound **4** (0.109 g, 0.229 mmol, 1 equivalent) was dissolved in anhydrous toluene (5 mL) and cooled to -78 °C. DIBAL-H (229 µL, 0.229 mmol, 1 equivalent) was added, dropwise. After ∼1 minute, took a TLC (4:1 hexanes:ethyl acetate), which indicated the production of product with an Rf of 0.3 and the consumption of **4**. The crude residue (**17,** 0.064 g, 59 %) was used without further purification.

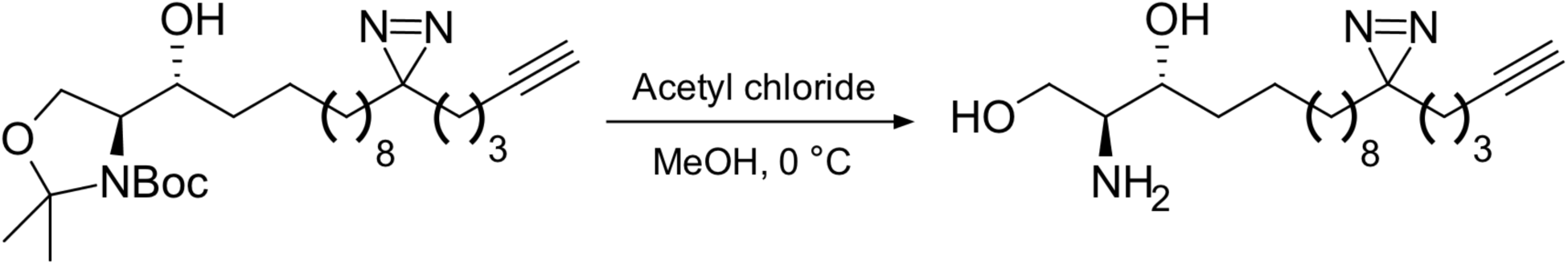

**(2R, 3S)-2-amino-13-[3-pent-4-yn-1-yl)-3H-diazirin-3-yl]tridecane-1,3-diol (5)** Compound **17** (64 mg, 0.134 mmol, 1 equivalent) was dissolved in methanol (5 mL) and cooled to 0 °C. Acetyl chloride (about 400 µL, until the pH reached 1) was added. The reaction proceeded overnight, until TLC (50:5:1 DCM : methanol : ammonium hydroxide, potassium permanganate staining) indicated production of product (Rf = 0.3; Rf of **17** in this system = 0.95). The methanol was evaporated and then 10 mL water and 10 mL DCM were added, along with 1M NaOH, until pH > 10 (∼ 2 mL). The aqueous layer was extracted 3 x 10 mL DCM and washed 2x with brine, dried over magnesium sulfate, and concentrated under reduced pressure. The crude residue was purified by flash chromatography, same solved system as above, producing pure **5** (25 mg, 55% yield). ^1^H NMR (400 MHz, MeOD) δ = 3.575 (m, 4H), δ = 3.122 (m, 1H), δ = 3.005 (m, 1H), δ = 2.210 (t, J = 1.4, 1H), δ = 2.156 (td, J = 6.8, J = 1.4, 2H), δ = 1.497 (m, 4H), δ = 1.301 (m, 20H), δ = 1.10 (m, 2H); [M + H]+, calcd. for C_19_H_35_N_3_O_2_, 338.2810, found 338.4.

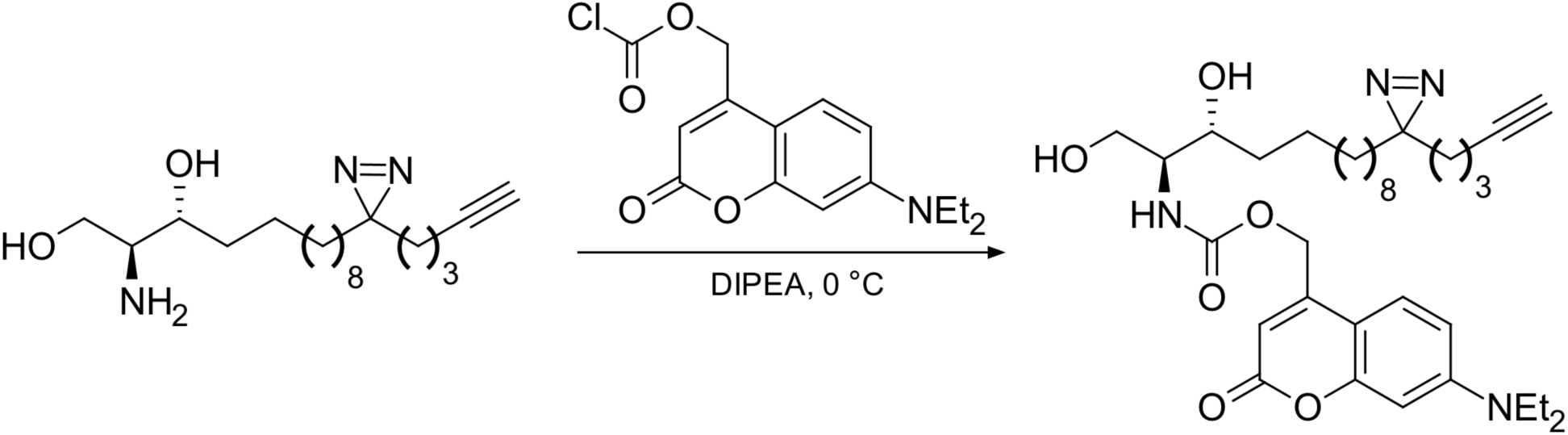

**[7-(diethylamino)-2-oxo-2H-chromen-4-yl]methyl N-[(2S, 3R)-1,3-dihydroxy-13[3-(pent-4- yn-1-yl)-3H-diazirin-3-7l]tridecan-2-yl] carbamate (TF-Spa, 1)** Coumarin chloroformate was produced in situ as previously described. 7-diethylamino-4-hydroxycoumarin (40 mg, 0.1618 mmol, 1 equivalent) was dissolved in 4 mL THF and cooled to 0 °C. DIPEA (84.5 µL, 0.4852 mmol, 3 equivalents). In a second flask, phosgene (1 mL [15 wt% in toluene], 1.618 mmol, 10 equivalents) was dissolved in 4 mL THF and cooled to 0 °C. The coumarin solution was added to the phosgene, dropwise, and the reaction was stirred at 0 °C for 2 hours. TLC (1:1 hexanes : ethyl acetate) revealed formation of product (Rf = 0.9) and the reaction was worked up: 10 mL water was added, and the reaction was extraction 3x10 mL with ethyl acetate and washed 1x20 mL with brine, dried over magnesium sulfate, and concentrated under reduced pressure. The crude residue was used without purification. Compound **5** (25 mg, 0.075 mmol, 1 equivalent) was dissolved in 3 mL dry THF and cooled to 0 °C. The crude coumarin chloroformate was dissolved in 2 mL dry THF and added dropwise to compound 19. The reaction was allowed to warm to room temperature and proceeded overnight, when TLC (50 : 5 : 1 DCM : methanol : ammonium hydroxide) revealed the complete consumption of 19 (Rf = 0.3, visible only by potassium permanganate staining) and the production of product (Rf = 0.5, visible by potassium permanganate staining and irradation with 375 nm light). 30 mL citric acid was added and the reaction was extracted 3 x 30 mL ethyl acetate and washed 1x10% sodium bicarbonate and 1xbrine, and dried over magnesium sulfate and concentrated under reduced pressure. The residue was purified twice by flash chromatography (once with 1:1 hexanes : ethyl acetate to remove residual 7-diethylamine-4-hydroxycoumarin and once with 10:1 DCM : methanol), and finally by HPLC, with a gradient of acetonitrile in water, 20% to 100% acetonitrile over 35 minutes, producing **TF-Spa, 1,** as a bright yellow oil (13.7 mg, 30% yield). ^1^H NMR (400 MHz, MeOD) δ = 7.365 (d, J = 8.8 1H), δ = 6.649 (s, 1H), 6.754 (m, 1H), 6.209 (s, 1H), 5.265 (m, 3H), 4.089 (dd, J = 11.6, J = 2.8, 1H), 3.994 (t, J= 5.6, 1H), 3.865 (m, 2H), 3.708 (m, 1H), 3.633 (m, 1H), 4.57 (q, J = 6.8, 4H), 2.179 (td, J = 6.6, J = 2.8, 2H), 1.964 (t, J = 6.6, J = 2.8, 1H), 1.581 (m, 4H), 1.507 (m, 3H), 1.362 (m, 9H), 1.257 (m, 25H), 1.091 (m, 3H); ^13^C NMR (400 MHz, CDCl_3_) δ = 124.634, 120.963, 83.528, 68.906, 61.887, 34.606, 34.323, 32.894, 31.885, 29.749, 29.548, 29.530, 29.502, 29.461, 29.400, 29.219, 28.517, 25.970, 25.607, 23.862, 22.806, 180.099, 18.011, 12.215, 4.661, 1.066; [M + H]+, calcd. for C_34_H_50_N_4_O_6_, 611.3810, found 611.6.

### Cell-based experiments

#### Calcium imaging and sphingoid base uncaging

HeLa cells were seeded to 70% density in an 8-well Labtek dish. Each well was loaded with 100 µL of a 5 µM Fluo-4 AM (Molecular Probes) solution in imaging buffer (20 mM HEPES, 115 mM NaCl, 1.2 mM MgCl_2_, 1.2 mM K_2_HPO_4_, 1.8 mM CaCl_2_, 10 mM glucose), and caged lipids were added to a final concentration of 2 µM. Fluorescence images of the Fluo-4 treated cells were captured on an inverted dual scanner confocal laser scanning microscope (Olympus Fluoview 1200) with a 63x oil objective using excitation at 488 nm for Fluo-4 imaging and simultaneous excitation at 375 nm for uncaging.

#### Analysis of trifuncitonal lipids by thin-layer chromatography

##### Probe labeling

Huh7 cells were seeded in 6 cm dishes and grown to confluence. Dilutions of trifunctional probe were made in complete media for the respective cell line; 2 µM sphingoid bases were used. 1 mL probe dilution was added to each plate and allowed to sit on cells for 30 min at 37°C prior to uncaging. Trifunctional probes were uncaged and photocrosslinked using two lamps: for crosslinking, a lamp with narrow band 350nm (NailStar 36 W UV lamp, Amazon, ASIN B00R4M0TI0); and for uncaging a lamp with narrow band 400nm (NailStar LED lamp, Amazon, ASIN B01286DTFQ). After removing the probe media and replacing it with normal complete media, dishes were exposed to 400 nm light to uncage the probe, and returned to the incubator for varying amounts of time to allow for metabolism. *Lipid extraction* Lipids were extracted with a modified Bligh-Dyer extraction. Briefly, dishes were rinsed with PBS, and 1 mL of a 2 : 0.8 methanol:water mixture was added. Cells were scraped into this mixture and moved to a glass tube, to which 1 mL of chloroform was added. Tubes were vortexed to mix, and the layers were allowed to separate at 4 °C for 1 hour. Tubes were centrifuged (3,000 x g for 10 minutes) to ensure complete separation of layers, and then the chloroform layer was removed to a fresh tube. Fresh chloroform was added to the aqueous layer and the extraction was repeated; the chloroform layers were combined and then dried under a stream of nitrogen. *Click labeling* Extracted lipids were re-dissolved in 10 µL chloroform, and 40 µL of a click mix (5 µL each of 1 mM TBTA, 10 mM copper sulfate, 10 mM sodium ascorbate, and 20 µM CalFluor 647 azide, and 20 µL ethanol) was added. The reaction was allowed to proceed for 1 hour in the dark, and then extracts were once more dried under a stream of nitrogen. *Plating and running TLC* Extracted lipids were re-dissolved in 10 µL and plated on 10x10 cm HPTLC silica 60 glass plates without F254 fluorophore. Lipids were resolved by a 2-step system: first using chloroform/methanol/ saturated aqueous ammonium hydroxide 65:25:4 for 6 cm, then drying, and finally using hexane/ethyl acetate 1:1 for 9 cm. Fluorescently labeled lipids were visualized using a Sapphire molecular imager.

#### Precursor ion scanning to identify metabolites of trifunctional lipids

Probe labeling and lipid extraction were performed as described above. Huh7 cells were treated with each probe, exposed to 400 nm light for uncaging, and metabolism was allowed to proceed for 1 hour prior to lipid extraction. Dried lipid extracts were resuspended in 200 µL of system A (5 % water, 95 % acetonitrile, 1 mM ammonium acetate, pH = 8.2) and separated by HPLC with the following parameters (system B = 50 % water, 50 % acetonitrile, 1 mM ammonium acetate, pH =8.2; column = 00D-437-BD, Luna 3 µm NH_2_ 100 Å LC column 100 x 2 mm, S/N H18-106425; B/ N 5377-0020):

| Time (min) | %A | %B | Flow rate (mL/min) |
| --- | --- | --- | --- |
| 0 | 99.9 | 0.1 | 0.3 |
| 6 | 94 | 6 | 0.3 |
| 10 | 75 | 25 | 0.3 |
| 13 | 0 | 100 | 0.3 |
| 13.5 | 0 | 100 | 0.5 |
| 18.6 | 99.9 | 0.1 | 0.5 |
| 18.7 | 99.9 | 0.1 | 0.5 |
| 23 | 99.9 | 0.1 | 0.5 |
| 23.5 | 99.9 | 0.1 | 0.7 |
| 24 | Stop |  |  |

Precursor ion scanning was used to identify precursors of phosphocholine in the positive mode, eluting between 3 and 5 minutes; collision induced dissociation was induced with an ESI ion source; ion spray voltage of 4000-4500 V; collision energy of 40; and declustering potential of 100-120.

#### Subcellular visualization of lipids by confocal microscopy

##### Probe labeling

Huh7 cells were seeded either in glass-bottomed 96-well plates, glass-bottomed 24-well plates, or on coverslips in 24-well plates, and grown to about 70% confluence. Dilutions of trifunctional probe were made in complete media for the respective cell line; 2 µM trifunctional sphingoid bases were used. 70 µL probe dilution (for 96-well plates; 200 µL for 24-well plates) was added to each plate and allowed to sit on cells for 30 min at 37 °C prior to uncaging. Dishes were exposed to 400 nm light to uncage the probe, and returned to the incubator for varying amounts of time to allow for metabolism, then exposed to 350 nm to photo-crosslink the probe, and immediately fixed by washing twice with PBS, then left in methanol for 20 min. *Click labeling* Cells were washed three times with PBS to remove organic solvent, and then 200 µL (for 24-well plates) or 70 µL (for 96-well plates) of a click mix were added (1 mM copper sulfate, 1 mM sodium ascorbate, 100 µM TBTA, 2 µM A647 picolyl azide, in PBS). The reaction was allowed to proceed for 1 h in the dark. *Antibody staining* Click mix was removed, cells were washed twice with PBS, and blocking buffer (2% BSA, 0.1% Triton-X-100 in PBS) was added. Cells were blocked for 1 h before the addition of primary antibodies. All primary antibodies (anti-Giantin, anti-PDI, and anti-Lamp1, catalogue numbers listed above) were diluted 1:250 in blocking buffer, and left on cells, with rocking, overnight at 4 °C. The next day, primary antibodies were removed, cells were washed three times with PBS, and fluorescently tagged secondary antibodies were added, either A488 anti-rabbit or A555 anti-mouse, 1:500 dilution in blocking buffer, for 1 h at room temperature, with rocking. Secondary antibodies were removed, cells were washed three times with PBS, and DAPI (1:1000) was added for ten minutes. Cells were imaged within a week after staining, on a Zeiss LSM 980 Laser-Scanning 4-channel confocal microscope with Airyscan.2^27^. *Image analysis* Pearson’s correlation coefficients between the lipid signal and the signal for each organelle marker were calculated using a CellProfiler pipeline^28^. Individual cells were selected based on regions of intensity of the lipid signal and coefficients were calculated within each cell.

#### Isolation and identification of protein-lipid complexes

##### Probe labeling

Huh7 cells were seeded in 10 cm dishes and grown to confluency. Dilutions of trifunctional probe were made in complete media for the respective cell line; 2 µM sphingoid bases were used. 3 mL of probe dilution was added to each plate and allowed to sit on cells for 30 min at 37 °C prior to uncaging. Cells treated with sphingoid bases were subjected to photo-crosslinking 15 min after uncaging; cells treated with fatty acids were subjected to photo-crosslinking 1 h after uncaging. After photo-crosslinking, cells were washed three times with PBS and scraped into 2 mL of ice-cold PBS. Cells were pelleted by centrifugation (1,000 x g for 5 minutes), supernatant was decanted, and cells were resuspended in 500 µL PBS.

##### Sample preparation for proteomics

Cells were lysed by probe sonication, on ice, in three 15 sec bursts. Lysates were subjected to a click reaction with picolyl azide agarose beads: 200 µL azide beads were washed once in DI water, then added to the cell lysate, along with copper sulfate (1mM, final concentration), sodium ascorbate (1 mM, final concentration), and TBTA (100 µM, final concentration). Samples were rotated at room temperature for 1 h. Beads were spun down (1,000 x g for 2 min), transferred to 2 mL centrifuge columns, and washed extensively: 3x with PBS, 5x with bead wash buffer 1 (100 mM Tris-HCl, pH = 8.0, 250 mM NaCl, 5 mM EDTA, 1% SDS), 10x with bead wash buffer 2 (100 mM Tris-HCl, pH = 8.0, 8M urea). Beads were transferred from the column in PBS to a clean Eppendorf tube and spun down. Isolated proteins were reduced (by re-suspending them in 1 mM digestion buffer [100 mM Tris-HCl, pH = 8.0, 2 mM CaCl_2_, 10% ACN], adding DTT to 10 mM, and incubating at 42 °C for 30 minutes), alkylated (by spinning down beads and resuspending them in 1 mL 40 mM aqueous iodoacetamide and incubating them at room temperature in the dark for 30 minutes. Bead-bound proteins were then digested by spinning the beads down, adding 50 µL of digestion buffer and 1 µL of LC-MS grade trypsin, and shaken at 37 °C overnight. Peptides were then desalted on C18 columns and the eluent was frozen at -80 °C.

##### Identification of isolated proteins by LC-MS/MS

Dried peptides were shipped to the EMBL proteomics core facility where they were TMT-labeled using the TMT-16-plex system and analyzed by LC-MS/MS on an Orbitrap Fusion Lumos mass spectrometer (Thermo Scientific). Peptides were separated using an Ultimate 3000 nano RSLC system (Dionex) equipped with a trapping cartridge (Precolumn C18 PepMap100, 5 mm, 300 “m i.d., 5 “m, 100 Å) and an analytical column (Acclaim PepMap 100. 75 × 50 cm C18, 3 mm, 100 Å) connected to a nanospray-Flex ion source. The peptides were loaded onto the trap column at 30 µl per min using solvent A (0.1% formic acid) and eluted using a gradient from 2 to 80% Solvent B (0.1% formic acid in acetonitrile) over 2 h at 0.3 µl per min (all solvents were of LC-MS grade). The Orbitrap Fusion Lumos was operated in positive ion mode with a spray voltage of 2.2 kV and capillary temperature of 275 °C. Full scan MS spectra with a mass range of 375–1500 m/z were acquired in profile mode using a resolution of 120,000 with a maximum injection time of 50 ms, AGC operated in standard mode and a RF lens setting of 30%. Fragmentation was triggered for 3 s cycle time for peptide like features with charge states of 2–7 on the MS scan (data-dependent acquisition). Precursors were isolated using the quadrupole with a window of 0.7 m/z and fragmented with a normalized collision energy of 34%. Fragment mass spectra were acquired in profile mode and a resolution of 30,000. Maximum injection time was set to 94 ms and AGC target to custom. The dynamic exclusion was set to 60 s.

##### Data analysis

Acquired data were analyzed using IsobarQuant^29^ and Mascot V2.4 (Matrix Science) using a reverse UniProt FASTA Homo sapiens database (UP000005640) including common contaminants. The following modifications were taken into account: Carbamidomethyl (C, fixed), TMT16plex (K, fixed), Acetyl (N-term, variable), Oxidation (M, variable) and TMT16plex (N-term, variable). The mass error tolerance for full scan MS spectra was set to 10 ppm and for MS/MS spectra to 0.02 Da. A maximum of 2 missed cleavages were allowed. A minimum of 2 unique peptides with a peptide length of at least seven amino acids and a false discovery rate below 0.01 were required on the peptide and protein level^30^. Only proteins that were quantified with two unique peptide matches in both replicates were kept for the analysis. A variance stabilization normalization was performed on the log2 raw data^31^, and enrichments of proteins in the (+) UV condition over the (-) UV condition was calculated using LIMMA analysis^32^. A protein is considered a “hit” in a certain condition if the false discovery rate is smaller than 0.05 and the fold change is at least 2; a protein is considered a “candidate” if a the false discovery rate is smaller than 0.2 and the fold change is at least 1.5.

##### Analysis of GO terms

Gene Ontology^33^ terms for each hit or candidate protein were downloaded from UniProt. Similar terms were manually grouped (e.g., location terms such as “mitochondrial matrix” and “mitochondrial iron-sulfur complexes” were grouped as “mitochondria”) and terms for hit or candidate proteins for each probe were counted.

