## Supporting Information for "Trifunctional sphinganine: a new tool to dissect sphingolipid function"

**CHEMICAL SYNTHESIS**

*Chemicals and reagents for synthesis* All chemicals were purchased from commercial suppliers and were used without further purification. Solvents were of ACS chemical grade (Fisher Scientific) and were used without further purification. Commercially available starting reagents were used without further purification. Analytical thin-layer chromatography was performed on silica gel 60 F254 aluminum-backed plates (Millipore Sigma), and spots were visualized either by UV light (254 nm or 365 nm), or potassium permanganate staining (1.5 g $KMnO_4$, 10g $K_2CO_3$, and 1.25 mL 10% NaOH in 200 mL of water). Flash column chromatography was performed with manually packed columns using Thermo Scientific Chemical Silica gel (0.035-0.070mm, 60 Å). High pressure liquid chromatography (HPLC) was performed on a Varian Prostar 210 (Agilent) using Polars 5 C18-A columns (Analytical: 150 x 4.6 mm, 3 $\mu$m; Preparative: 150 x 21.2 mm, 5 $\mu$m). Mass spectra were obtained on a Advion Expressions CMS mass spectrometer. $^1$H NMR spetra were recorded on a Bruker DPX spectrometer at 400 MHz. Chemical shifts are reported as parts per million (ppm) relative to solvent references.

*Synthesis of TF-Spa (**1**)*

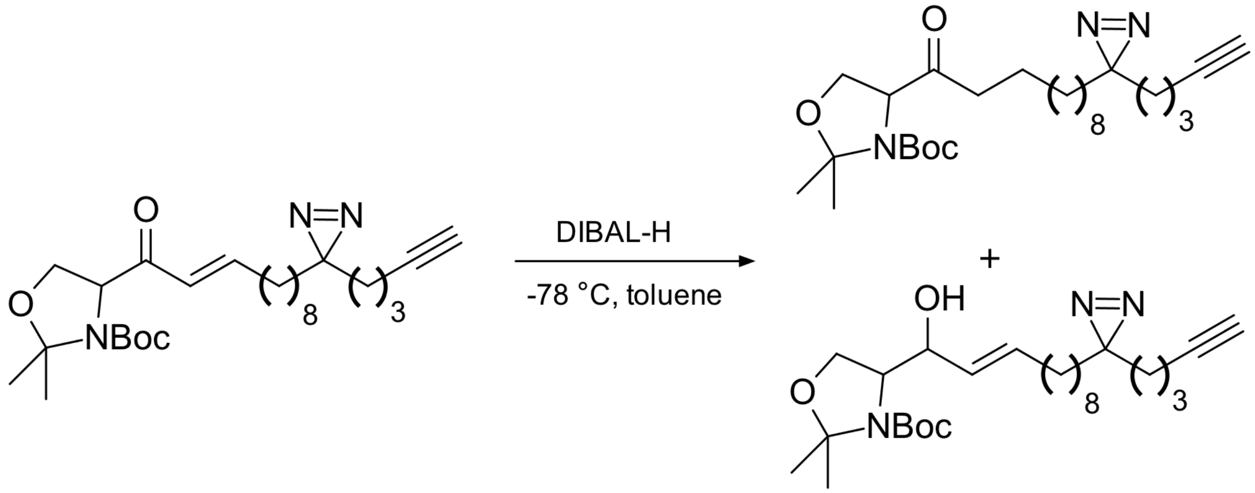


**Tert-butyl 2,2-dimethyl-4-{11-[pent-4-yn-1-yl)-3H-diazirin-3-yl]undecanoyl}-1,3-oxazolidine-3-carboxylate (4)** Compound **3** (0.50 g, 1.056 mmol, 1 equivalent) was dissolved in anhydrous toluene (15 mL) and cooled to -78 ˚C. DIBAL-H (1.1 mL, 1.056 mmol, 1 equivalent) was added, dropwise. After ~1 minute, took a TLC (4:1 hexanes:ethyl acetate, potassium permanganate visualization), which indicated the production of two products: the 1,4 reduction product, the ketone (**4**), with an Rf of 0.7 (Rf of **3** in this system is 0.65); and the 1,2 reduction product, the alpha-beta-unsaturated alcohol, with an Rf of 0.3. The 1,4 and the 1,2 reduction products were separated by flash chromatography (4:1 hexanes:ethyl acetate), producing ketone **4** as a pale yellow oil (0.109 g, 23 % yield), as well as the alpha-beta-unsaturated alcohol (0.100 g, 20% yield). $^1$H NMR (400 MHz, $CDCl_3$) δ = 4.472 (dd, J = 7.4, J = 3.2, 1H), δ = 4.325 (dd, J = 7.6, J = 2.8, 1H), δ = 4.155 (m, 2H), δ = 3.915 (m, 2H), δ = 2.494 (q, J = 8, 2H), δ = 2.178 (td, J = 6.8, J = 2.8, 2H), δ = 1.962 (t, J = 1.4, 1H), δ = 1.725 (s, 2H), δ = 1.666 (s, 2H), δ = 1.586 (s, 4H), δ = 1.513 (m, 9H), δ = 1.424 (s, 4H), δ = = 1.365 (m, 6H), δ = 1.25 (m, 12H), δ = 1.090 (m, 2H); $^{13}$C NMR (400 MHz, CD-

$Cl_3$) δ = 208.801, 208.360, 207.014, 152.485, 151.531, 138.083, 115.127, 95.247, 94.547, 83.568, 80.999, 80.663, 68.993, 68.310, 65.844, 65.486, 65.287, 39.184, 38.537, 33.029, 32.427, 31.947, 31.051, 29.819, 29.510, 29.430, 29.371, 28.557, 28.475, 26.324, 25.512, 24.967, 23.933, 23.205, 22.873, 18.077; [M + H]+, calcd. for $C_{29}H_{39}N_3O_4$, 476.3490, found 476.5.

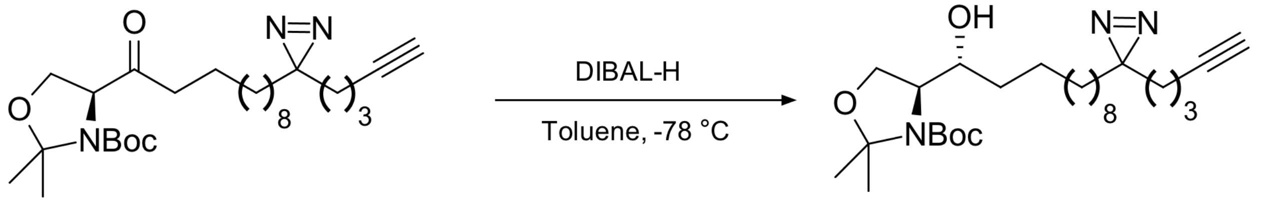


**(1R)-1-[3S)-5,5,-dimethyloxolan-3-yl]-11-[3-(pent-4-yn-1yl)-3H-diazirin-3-yl]undecan-1-ol (17)** Compound **4** (0.109 g, 0.229 mmol, 1 equivalent) was dissolved in anhydrous toluene (5 mL) and cooled to -78 ˚C. DIBAL-H (229 μL, 0.229 mmol, 1 equivalent) was added, dropwise. After ~1 minute, took a TLC (4:1 hexanes:ethyl acetate), which indicated the production of product with an Rf of 0.3 and the consumption of **4**. The crude residue (**17,** 0.064 g, 59 %) was used without further purification.

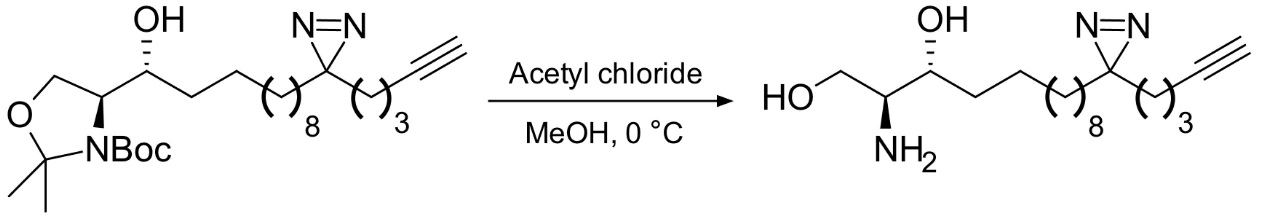


**(2R, 3S)-2-amino-13-[3-pent-4-yn-1-yl)-3H-diazirin-3-yl]tridecane-1,3-diol (5)** Compound **17** (64 mg, 0.134 mmol, 1 equivalent) was dissolved in methanol (5 mL) and cooled to 0 ˚C. Acetyl chloride (about 400 μL, until the pH reached 1) was added. The reaction proceeded overnight, until TLC (50:5:1 DCM : methanol : ammonium hydroxide, potassium permanganate staining) indicated production of product (Rf = 0.3; Rf of **17** in this system = 0.95). The methanol was evaporated and then 10 mL water and 10 mL DCM were added, along with 1M NaOH, until pH > 10 (~ 2 mL). The aqueous layer was extracted 3 x 10 mL DCM and washed 2x with brine, dried over magnesium sulfate, and concentrated under reduced pressure. The crude residue was purified by flash chromatography, same solved system as above, producing pure **5** (25 mg, 55% yield). $^1H$ NMR (400 MHz, MeOD) δ = 3.575 (m, 4H), δ = 3.122 (m, 1H), δ = 3.005 (m, 1H), δ = 2.210 (t, J = 1.4, 1H), δ = 2.156 (td, J = 6.8, J = 1.4, 2H), δ = 1.497 (m, 4H), δ = 1.301 (m, 20H), δ = 1.10 (m, 2H); [M + H]+, calcd. for $C_{19}H_{35}N_3O_2$, 338.2810, found 338.4.

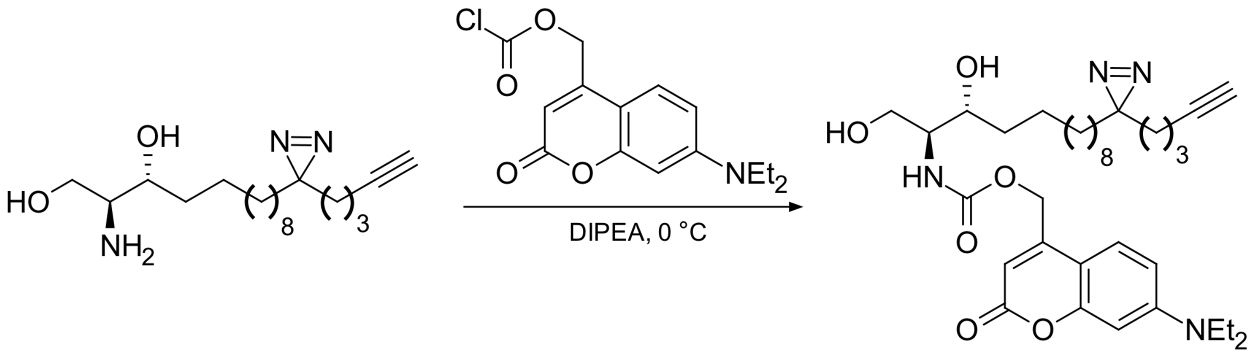


**[7-(diethylamino)-2-oxo-2H-chromen-4-yl]methyl N-[(2S, 3R)-1,3-dihydroxy-13[3-(pent-4-yn-1-yl)-3H-diazirin-3-7l]tridecan-2-yl] carbamate (TF-Spa, 1)** Coumarin chloroformate was produced in situ as previously described. 7-diethylamino-4-hydroxycoumarin (40 mg, 0.1618 mmol, 1 equivalent) was dissolved in 4 mL THF and cooled to 0 ˚C. DIPEA (84.5 $\mu$L, 0.4852 mmol, 3 equivalents). In a second flask, phosgene (1 mL [15 wt% in toluene], 1.618 mmol, 10 equivalents) was dissolved in 4 mL THF and cooled to 0 ˚C. The coumarin solution was added to the phosgene, dropwise, and the reaction was stirred at 0 ˚C for 2 hours. TLC (1:1 hexanes : ethyl acetate) revealed formation of product (Rf = 0.9) and the reaction was worked up: 10 mL water was added, and the reaction was extraction 3x10 mL with ethyl acetate and washed 1x20 mL with brine, dried over magnesium sulfate, and concentrated under reduced pressure. The crude residue was used without purification. Compound **5** (25 mg, 0.075 mmol, 1 equivalent) was dissolved in 3 mL dry THF and cooled to 0 ˚C. The crude coumarin chloroformate was dissolved in 2 mL dry THF and added dropwise to compound 19. The reaction was allowed to warm to room temperature and proceeded overnight, when TLC (50 : 5 : 1 DCM : methanol : ammonium hydroxide) revealed the complete consumption of 19 (Rf = 0.3, visible only by potassium permanganate staining) and the production of product (Rf = 0.5, visible by potassium permanganate staining and irradation with 375 nm light). 30 mL citric acid was added and the reaction was extracted 3 x 30 mL ethyl acetate and washed 1x10% sodium bicarbonate and 1xbrine, and dried over magnesium sulfate and concentrated under reduced pressure. The residue was purified twice by flash chromatography (once with 1:1 hexanes : ethyl acetate to remove residual 7-diethylamine-4-hydroxycoumarin and once with 10:1 DCM : methanol), and finally by HPLC, with a gradient of acetonitrile in water, 20% to 100% acetonitrile over 35 minutes, producing **TF-Spa, 1,** as a bright yellow oil (13.7 mg, 30% yield). $^1$H NMR (400 MHz, MeOD) δ = 7.365 (d, J = 8.8 1H), δ = 6.649 (s, 1H), 6.754 (m, 1H), 6.209 (s, 1H), 5.265 (m, 3H), 4.089 (dd, J = 11.6, J = 2.8, 1H), 3.994 (t, J= 5.6, 1H), 3.865 (m, 2H), 3.708 (m, 1H), 3.633 (m, 1H), 4.57 (q, J = 6.8, 4H), 2.179 (td, J = 6.6, J = 2.8, 2H), 1.964 (t, J = 6.6, J = 2.8, 1H), 1.581 (m, 4H), 1.507 (m, 3H), 1.362 (m, 9H), 1.257 (m, 25H), 1.091 (m, 3H); $^{13}$C NMR (400 MHz, $CDCl_3$) δ = 124.634, 120.963, 83.528, 68.906, 61.887, 34.606, 34.323, 32.894, 31.885, 29.749, 29.548, 29.530, 29.502, 29.461, 29.400, 29.219, 28.517, 25.970, 25.607, 23.862, 22.806, 180.099, 18.011, 12.215, 4.661, 1.066; [M + H] +, calcd. for $C_{34}H_{50}N_4O_6$, 611.3810, found 611.6.

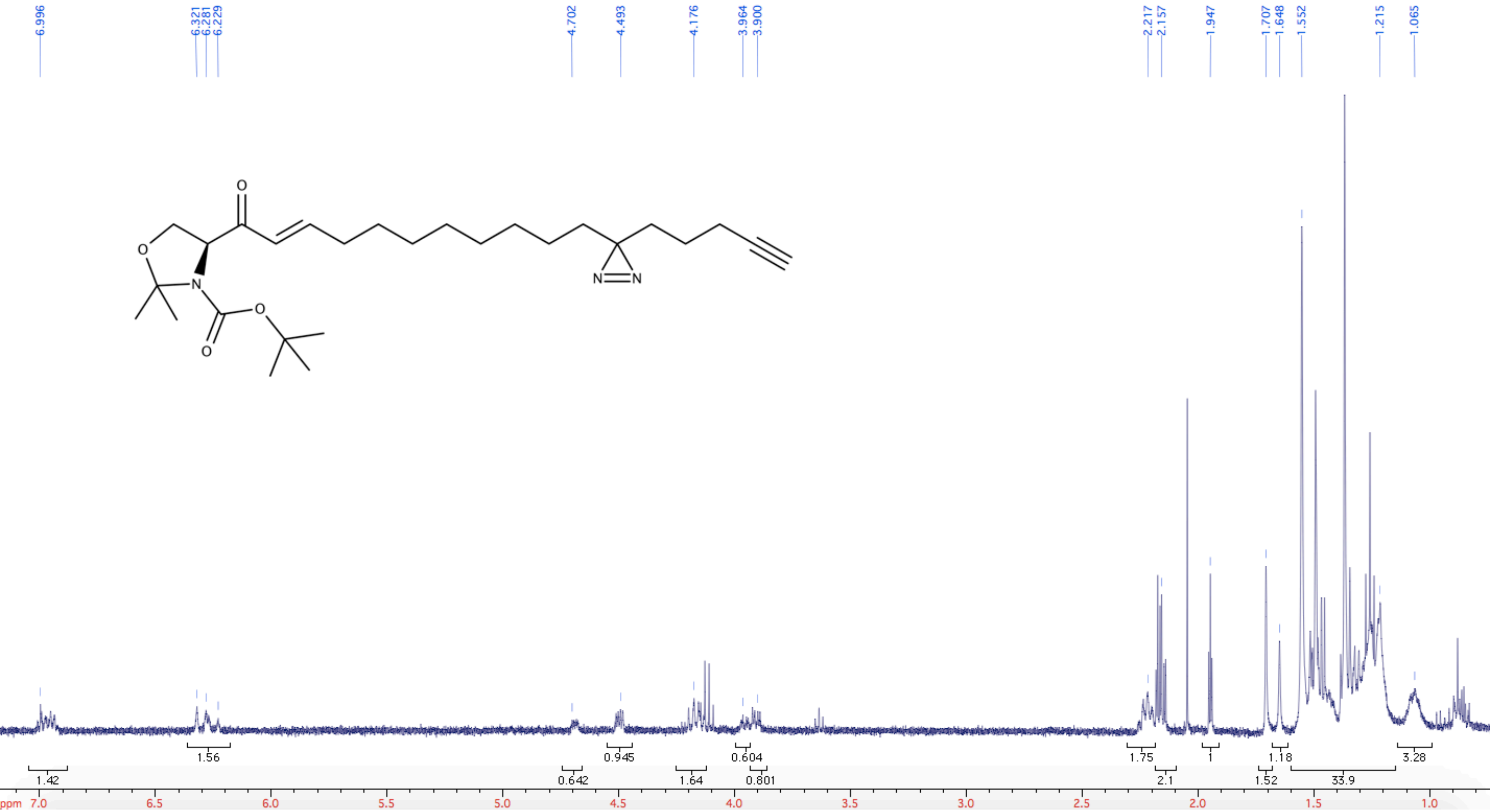
6.996
6.321
6.281
6.229
4.702
4.493
4.176
3.964
3.900
2.217
2.157
1.947
1.707
1.648
1.552
1.215
1.065
O
O
N
O
O
N
N
1.42
1.56
0.642
0.945
1.64
0.604
0.801
1.75
2.1
1
1.52
1.18
33.9
3.28
ppm
7.0
6.5
6.0
5.5
5.0
4.5
4.0
3.5
3.0
2.5
2.0
1.5
1.0

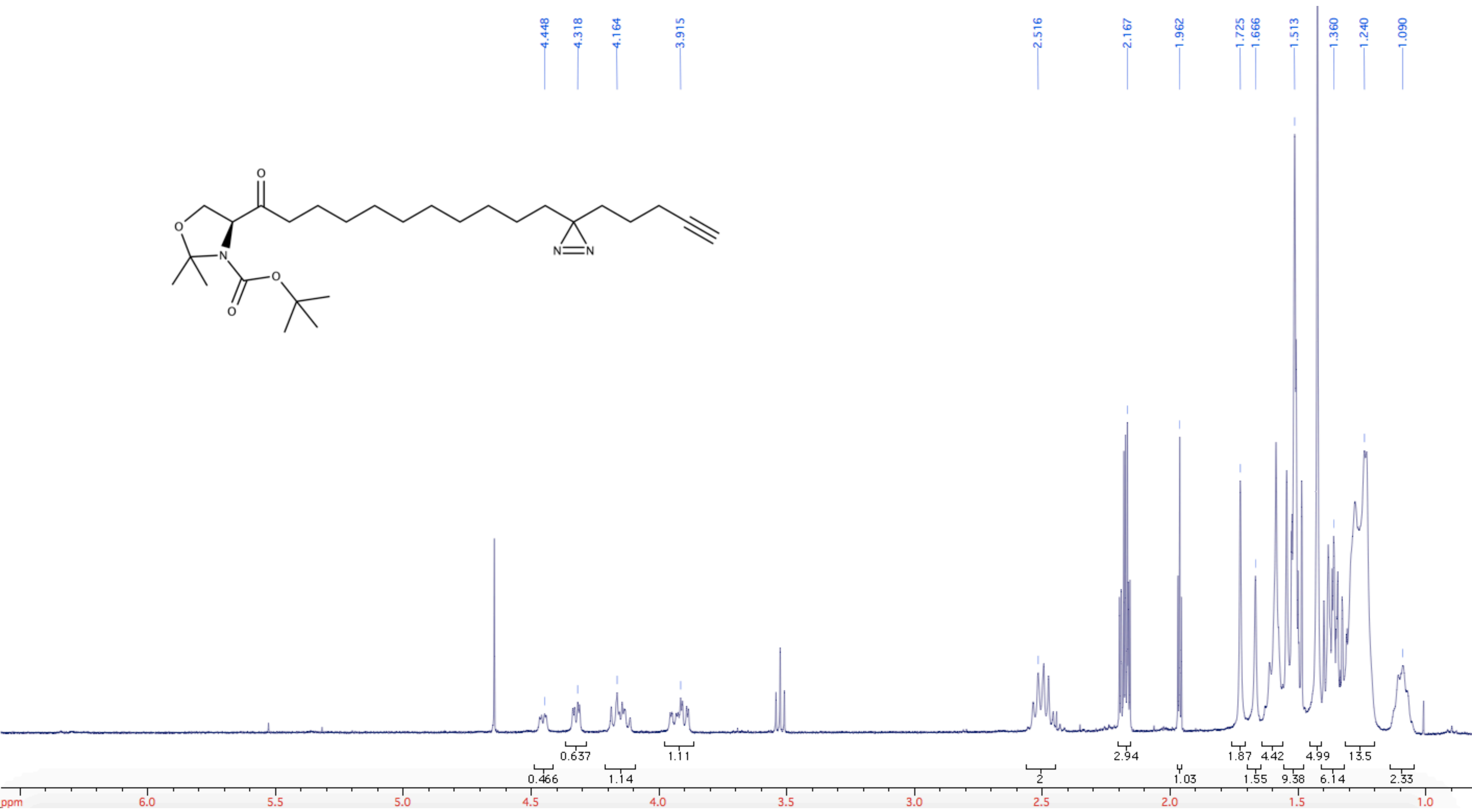
4.448
4.318
4.164
3.915
2.516
2.167
1.962
1.725
1.666
1.513
1.360
1.240
1.090
0.637
1.11
2.94
1.87
4.42
4.99
13.5
0.466
1.14
2
1.03
1.55
9.38
6.14
2.33
ppm
6.0
5.5
5.0
4.5
4.0
3.5
3.0
2.5
2.0
1.5
1.0
O
O
N
O
O
N
N

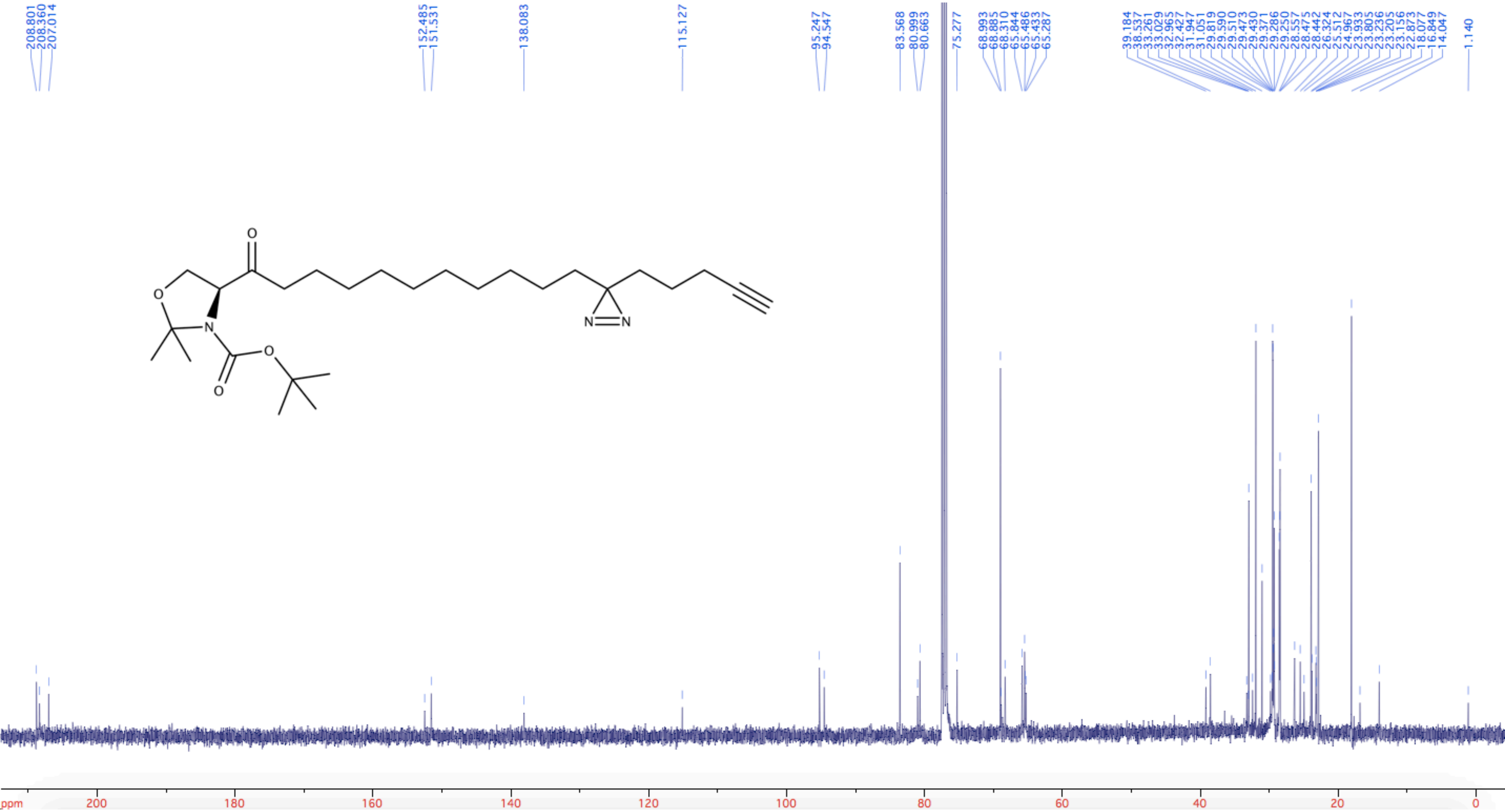

208.801
208.360
207.014
152.485
151.531
138.083
115.127
95.247
94.547
83.568
80.999
80.663
75.277
68.993
68.885
68.310
65.844
65.486
65.433
65.287
39.184
38.537
33.261
33.029
32.965
32.427
31.947
31.051
29.819
29.590
29.510
29.473
29.430
29.371
29.286
29.250
28.557
28.475
28.442
26.324
25.512
24.967
23.933
23.805
23.236
23.205
23.156
22.873
18.077
16.849
14.047
1.140
ppm
200
180
160
140
120
100
80
60
40
20
0

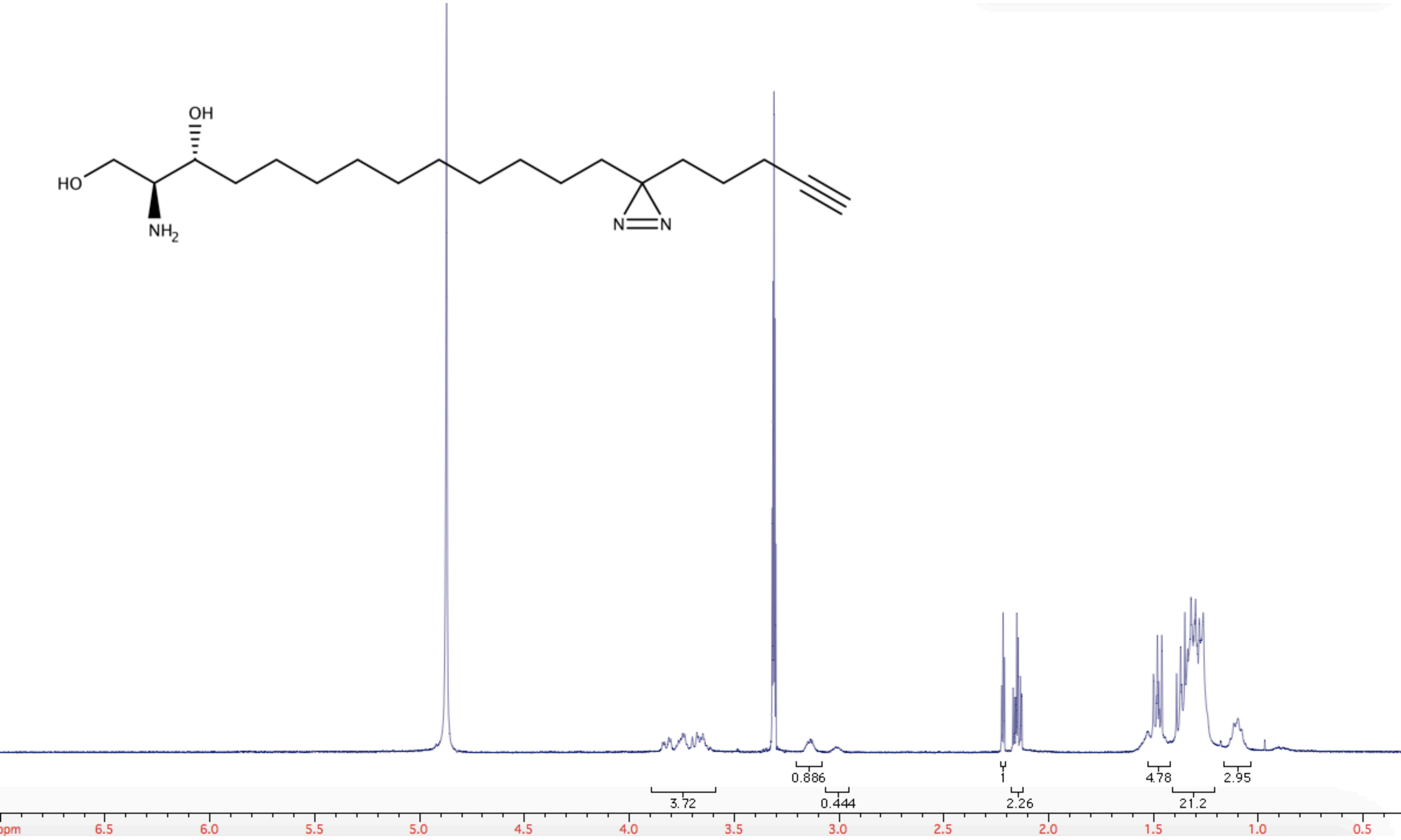
OH
HO
NH2
N
N
0.886
1
4.78
2.95
3.72
0.444
2.26
21.2
ppm
6.5
6.0
5.5
5.0
4.5
4.0
3.5
3.0
2.5
2.0
1.5
1.0
0.5

**TF-Spa**

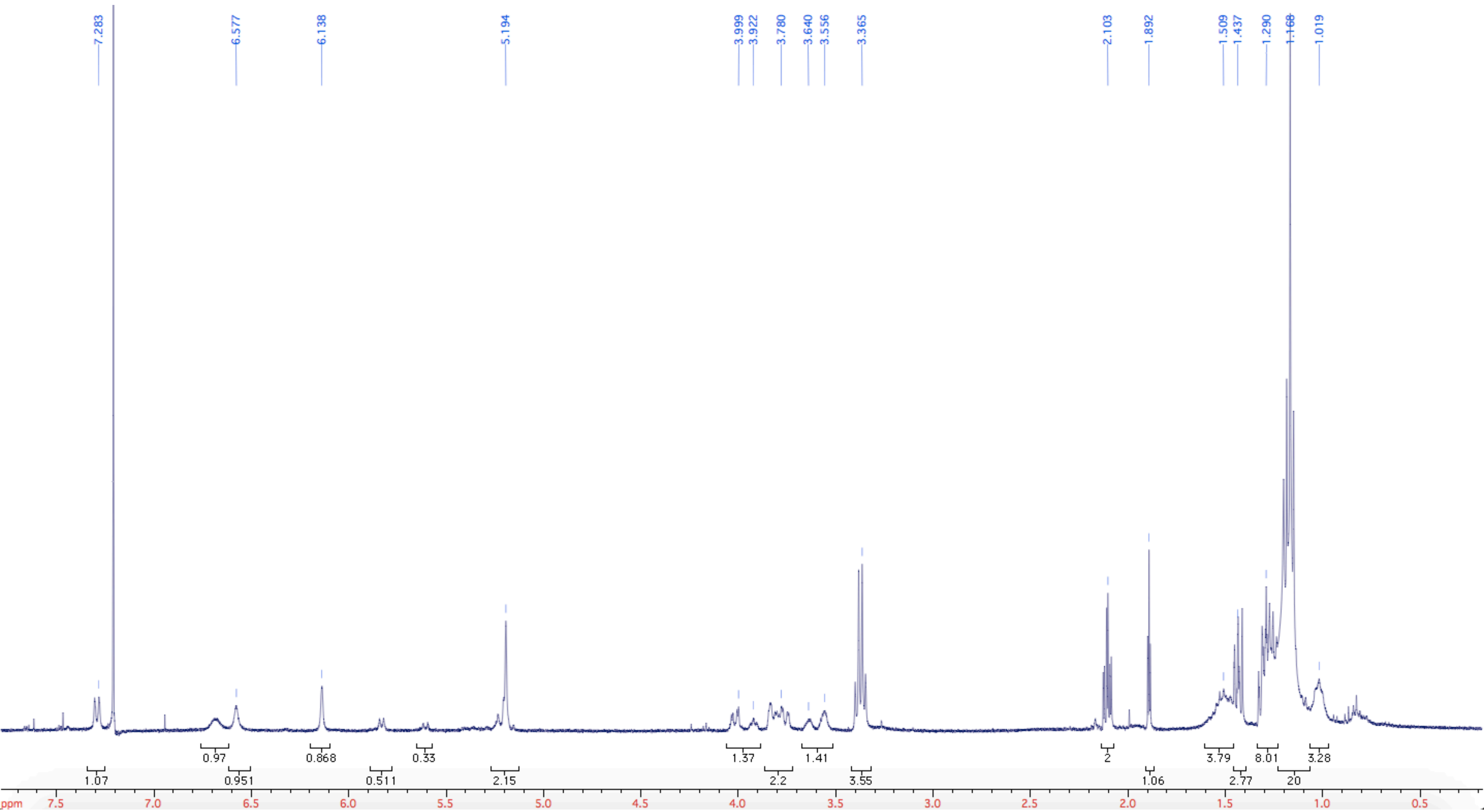
7.283
6.577
6.138
5.194
3.999
3.922
3.780
3.640
3.556
3.365
2.103
1.892
1.509
1.437
1.290
1.168
1.019
0.97
0.868
0.33
1.37
1.41
2
3.79
8.01
3.28
1.07
0.951
0.511
2.15
2.2
3.55
1.06
2.77
20
ppm
7.5
7.0
6.5
6.0
5.5
5.0
4.5
4.0
3.5
3.0
2.5
2.0
1.5
1.0
0.5

**TF-Spa**

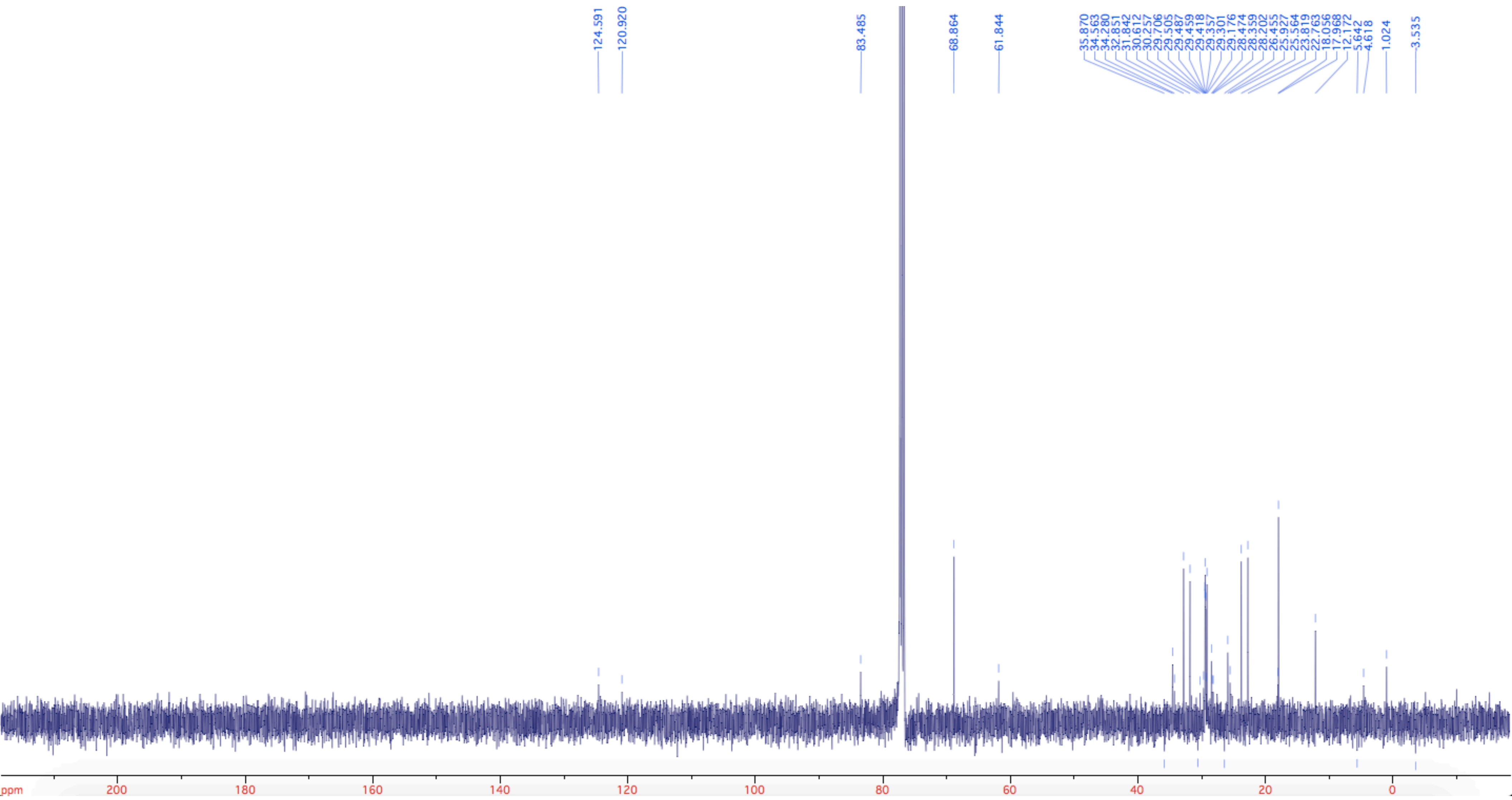
124.591
120.920
83.485
68.864
61.844
35.870
34.563
34.280
32.851
31.842
30.612
30.257
29.706
29.505
29.487
29.459
29.418
29.357
29.301
29.176
28.474
28.359
28.202
26.455
25.927
25.564
23.819
22.763
18.056
17.968
12.172
5.642
4.618
1.024
-3.535
ppm
200
180
160
140
120
100
80
60
40
20
0
